# Ligation bias is a major contributor to nonstoichiometric abundances of secondary siRNAs and impacts analyses of microRNAs

**DOI:** 10.1101/2020.09.14.296616

**Authors:** Patricia Baldrich, Saleh Tamim, Sandra Mathioni, Blake Meyers

**Affiliations:** Donald Danforth Plant Science Center, Saint Louis, Missouri, 63132, USA; Center for Bioinformatics and Computational Biology, University of Delaware, Newark, DE 19711, USA; Division of Plant Sciences, University of Missouri-Columbia, Columbia, MO 65211, USA

## Abstract

Plant small RNAs are a diverse and complex set of molecules, ranging in length from 21 to 24 nt, involved in a wide range of essential biological processes. High-throughput sequencing is used for the discovery and quantification of small RNAs. However, several biases can occur during the preparation of small RNA libraries, especially using low input RNA. We used two stages of maize anthers to evaluate the performance of seven commercially-available methods for small RNA library construction, using different RNA input amounts. We show that when working with plant material, library construction methods have differing capabilities to capture small RNAs, and that different library construction methods provide better results when applied to the detection of microRNAs, phasiRNAs, or tRNA-derived fragment. We also observed that ligation bias occurs at both ends of miRNAs and phasiRNAs, suggesting that the biased compositions observed in small RNA populations, including nonstoichiometric levels of phasiRNAs within a locus, may reflect a combination of biological and technical influences.

## INTRODUCTION

Small RNAs (sRNAs) are short non-coding molecules, ranging from 20 to 24 nt in length, with critical functions in all aspects of plant development and responses to the environment. High-throughput sequencing of sRNA libraries (sRNA-seq) is considered the most efficient way to accomplish sRNA discovery and quantify their abundances. Briefly, the process consists of a two-step DNA and RNA adapter ligation in which first the 3’ and then the 5’ adapters are ligated to the sRNAs, followed by retrotranscription and PCR amplification steps. These results in a dsDNA molecule comprised of an sRNA flanked by two adapter sequences (Figure 1). However, several issues can occur during library construction including (1) adapter ligation bias, (2) adapter dimerization, and (3) low RNA input amounts. Adapter ligation bias is due to sequence-derived differences in the efficiency of ligating one or both adapters(1). Several approaches have been developed to address this bias, such as the inclusion of degenerate ends in one or both adapters, the use of a single adapter for both 3’ and 5’ ligation, (taking advantage of intramolecular ligation efficiency), and even complete removal of the adapter ligation step by substituting sRNA polyadenylation. The second major issue during library preparation is adapter dimer formation. This is caused by the generation of an adapter-adapter molecule, with no sRNA insert, and it results in a loss of sequencing depth. This can be avoided by removing excess 3’ adapter molecules after the first ligation and/or performing a library size selection after PCR amplification, using beads or polyacrylamide gels. The third major issue is the use of low amounts of RNA as starting material, often driven by the necessity of doing single cell and low input transcriptomics to study multicellular organisms with a high heterogeneity and complexity. This can result in inefficient adapter ligation and a low complexity library.

**Figure 1:**
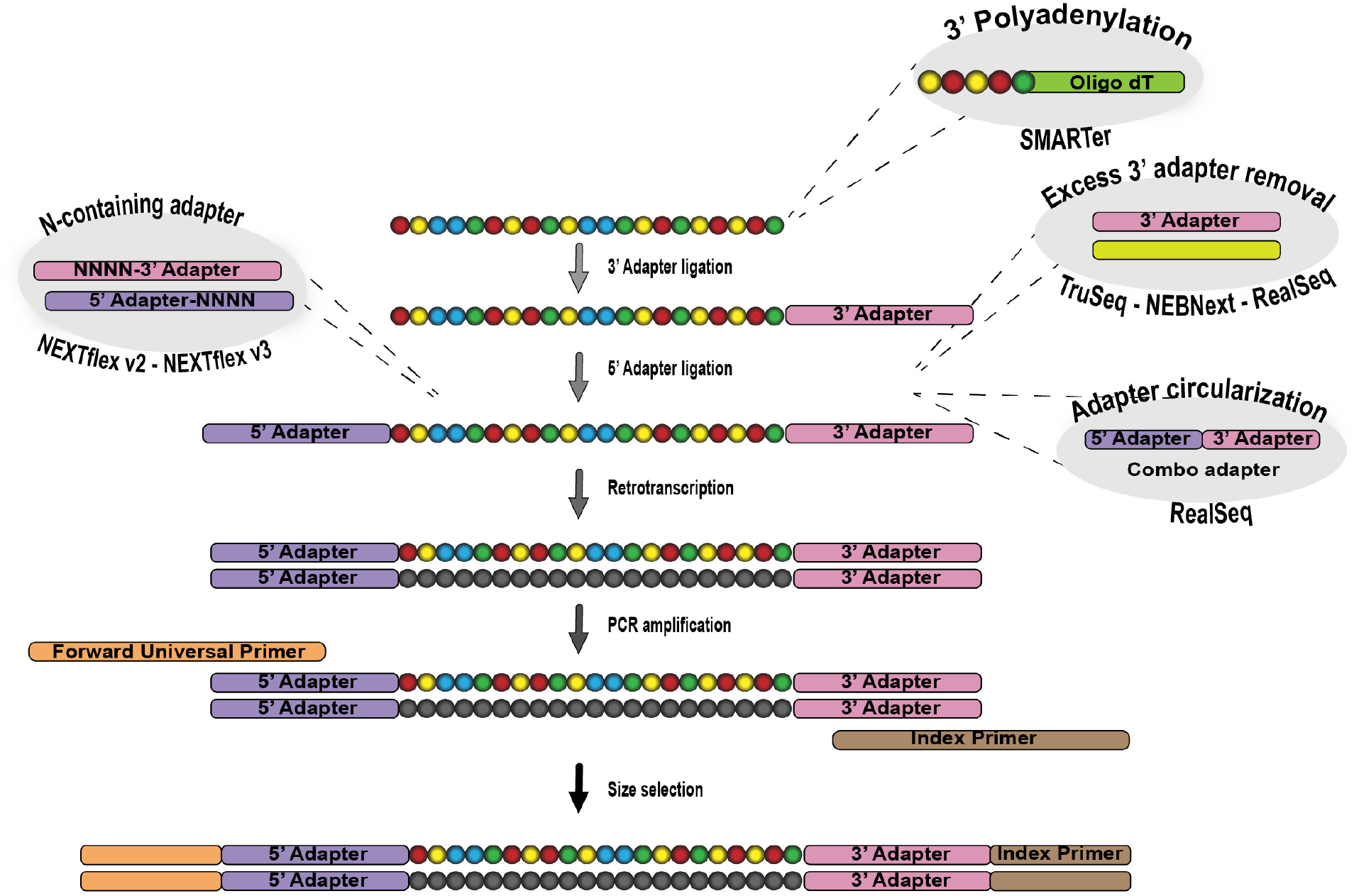
Workflow overview of the small RNA library preparation process. The general workflow is displayed as a vertical linear representation. Each step that is specific to a kit is displayed as a bubble.

In the last few years, several studies have focused on determining the most reliable, commercial methods for sRNA quantification (2–6). However, although informative, none of these studies used plant samples as input material. In this study, we focus on plant biological samples that are highly complex and include several different types of sRNAs. Based on their biogenesis pathways, plant sRNAs are divided into two main categories: microRNAs (miRNAs) and short-interfering RNAs (siRNAs). miRNAs derive from a self-complementary, single-stranded RNA, while siRNAs derive from an RNA molecule made double-stranded by an RNA-dependent RNA polymerase (7–9). siRNAs are further divided into several categories, with phased siRNAs (phasiRNAs) arguably the most-studied subset and recently shown to be essential elements in plant reproduction (10–12). A second type of siRNAs, heterochromatic siRNAs, comprise the majority of sRNAs in some species and are mainly involved in silencing of repetitive elements.

Maize anthers, the male reproductive organ, are an optimal source of sRNAs for this type of analysis as they produce substantial quantities of all of these sRNA classes. They are also one of the richest sources of phasiRNAs, including both 21- and 24-nt classes, shown to be important for the correct regulation of anther development. The development of maize anthers has been well characterized and thus it is an ideal model for studies of reproductive sRNAs (10, 13). Maize anthers have a unique pattern of sRNA expression; during the premeiotic stage, 21-nt phasiRNAs accumulate, triggered by miR2118. The later, meiotic anthers are enriched in 24-nt phasiRNAs, triggered by miR2275.

In this study, we used two different tissues from premeiotic (PMA) and meiotic (MA) maize anthers to evaluate the performance of seven different commercially available sRNA-seq methods of library construction.

## MATERIAL AND METHODS

### Biological material and RNA extraction

Anther samples were collected from 4- to 5-week-old W23 maize plants grown controlled conditions in greenhouses with temperatures of 28°C/22°C, relative humidity of 50%/60%, and a photoperiod of 16 h/8 h (day/night). Two developmental stages were dissected, the premeiotic anther stage consisting of spikelets with 0.2 to 0.7 mm anthers, and the meiotic anther stage consisting of 1.5 mm anthers. Samples were flash frozen in liquid nitrogen and kept at −80°C until further processing.

Total RNA was isolated using the Plant RNA Reagent (Thermo Fisher, cat #12322012). Briefly, samples were ground in a liquid nitrogen-frozen mortar and pestle. Then, the cold mortar with the powdered tissue was transferred to a chemical safe hood and 1 ml of Plant RNA Reagent was added directly in the mortar. The sample was ground with the reagent for another 20 sec and then collected with a 1ml micropipette and transferred to a 2 ml nuclease-free microcentrifuge tube. The remaining steps for the RNA extraction were followed as described previously (14). More details are available in Supplemental table 1 and Supplemental methods.

### Small RNA library generation and sequencing

All sRNA libraries were generated following the manufacturers’ instructions. A summarized workflow can be seen in Figure 1. Additionally, we performed a final step of size selection using 6% polyacrylamide gels as described in Mathioni *et al*., 2017 (14). All samples were sequenced using Illumina HiSeq 2500 at the University of Delaware DNA Sequencing & Genotyping Center at the Delaware Biotechnology Institute. Additional details can be found in Supplemental Methods.

#### Data analysis and visualization

The sRNA sequencing libraries were trimmed for adaptors using Trimmomatic v0.32 (15) with the following options -phred33; -threads 10; ILLUMINACLIP:4:30:10; LEADING:3; TRAILING:3; SLIDINGWINDOW:4:10; MINLEN: 18. Sequence quality was assessed using FastQC (http://www.bioinformatics.babraham.ac.uk/projects/fastqc/). Clean reads were aligned to the B73 Maize genome version 4 (16) using the software Bowtie2 (Langmead and Salzberg, 2012). For miRNA analyses, the latest version of miRBase (v22; (17)) was used. For phasiRNAs, the *PHAS* loci published in Zhai et al. (2015) were used. The genome coordinates for the most abundant *PHAS* loci are included in Supplemental table 2. All alignments for sRNA counts were performed using Bowtie2 (18). Differential accumulation of sRNAs was calculated using DESeq2 software (19). Data visualization was performed using R studio (20). In general, ggplot2 (21) and gridExtra (22) packages were used for plotting, and RColorBrewer was used for color schemes. Correlation plots were generated using the corrplot package (23). UpSet plots were generated using UpSetR pakage (24).

## RESULTS

### Library composition varies by method for distinct subsets of small RNAs

We isolated RNA from two different pools of maize anthers at premeiotic (PMA) and meiotic (MA) stages (see methods) and we generated small RNA (sRNA) libraries using different commercial kits. We utilized seven kits from six vendors, named as follows, with full details in the methods section: TruSeq, NEBNext, NEXTFlex v2, NEXTFlex v3, RealSeq, SMARTer, and TriLink. We performed three technical replicates of each library, for both PMA and MA stages, and for each input amount of RNA (see Table 1 below). All libraries were sequenced using the same Illumina HiSeq 2500 instrument, and to a similar depth (Supplemental Figure 1 – left boxplot). All the libraries were independently processed and mapped to the B73 maize genome v4 (25). We obtained similar mapping percentages for all libraries (Supplemental Figure 1); all subsequent analyses utilized these mapped reads.

**Table 1:**
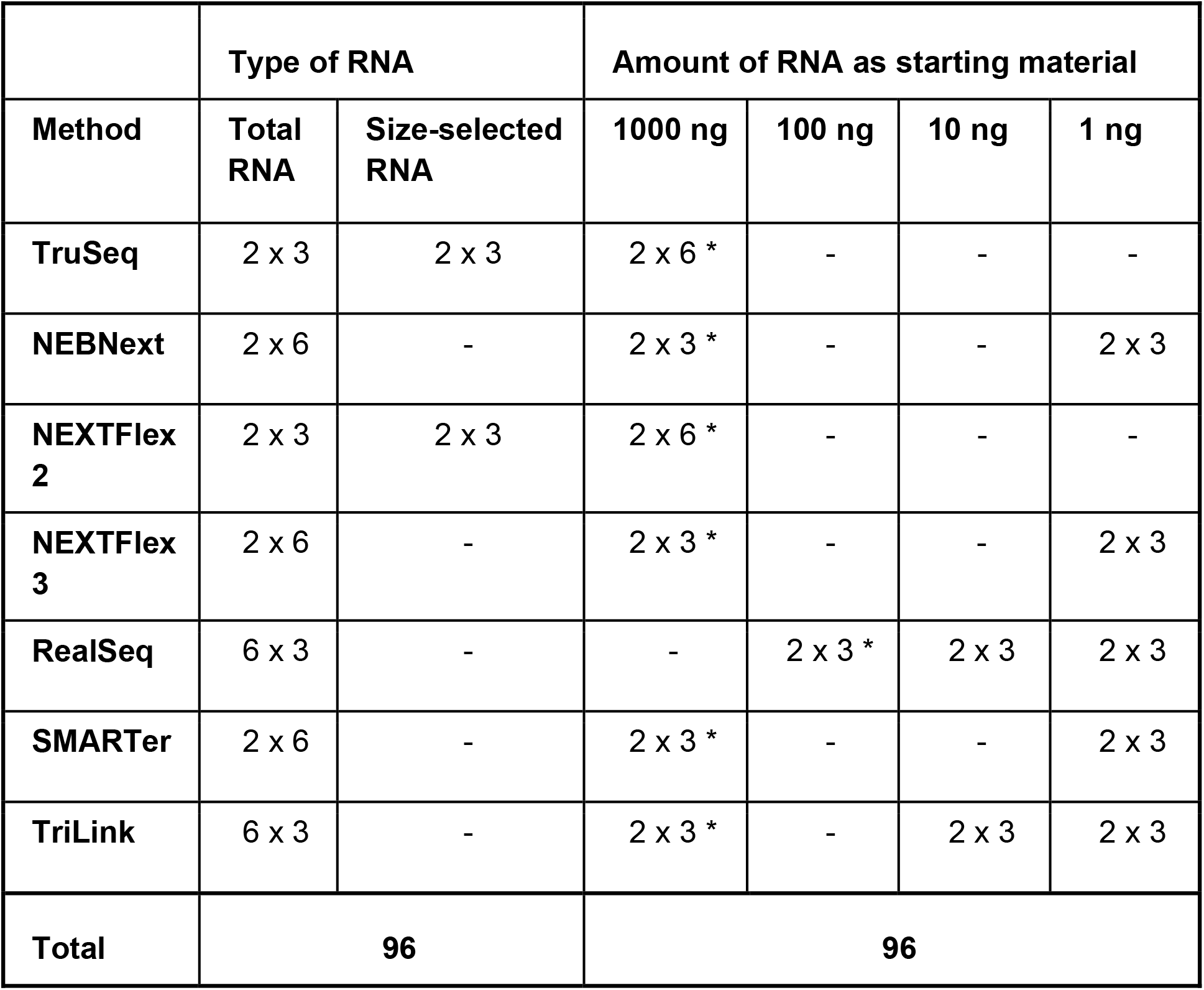
Summary of libraries included in this study. * indicates the recommended amount from the manufacturer. Three technical replicates were always used for each condition. Libraries were constructed in parallel for premeiotic anthers and meiotic anthers.

Size distribution is often used to analyze and evaluate the quality of sRNA libraries. We assessed two different components for each size: the total read abundance and the number of distinct counts. The latter component collapses together all the reads that have the same sequence and counts them as one, thus reflecting the diversity of reads. sRNAs are 20 to 24 nucleotide (nt) in size and their accumulation varies depending on tissue identity and species. Maize anthers are known to accumulate a high number of 24 nt sRNAs (26). When comparing the different library construction methods, we observed that TruSeq, NEBNext, NextfFlex and RealSeq generated a similar size distribution, with a predominant peak at 24 nt and a secondary peak at 21 nt, while SMARTer and TriLink generated a flatter distribution (Figure 2). This is due to the high abundance of structural RNAs present in these methods (see below). In PMA anthers, TruSeq and RealSeq methods demonstrated the highest peak at 24 nt, while both NEXTFlex methods yielded higher peaks at 21 and 22 nt. However, in MA anthers, we observed the opposite; both NEXTFlex methods demonstrated a higher peak at 24 nt and smaller peaks at 22 and 23 nt. This suggests that each kit yields a specific profile and captures diversity in a variable manner, potentially dependent on intrinsic properties of the sRNAs.

**Figure 2:**
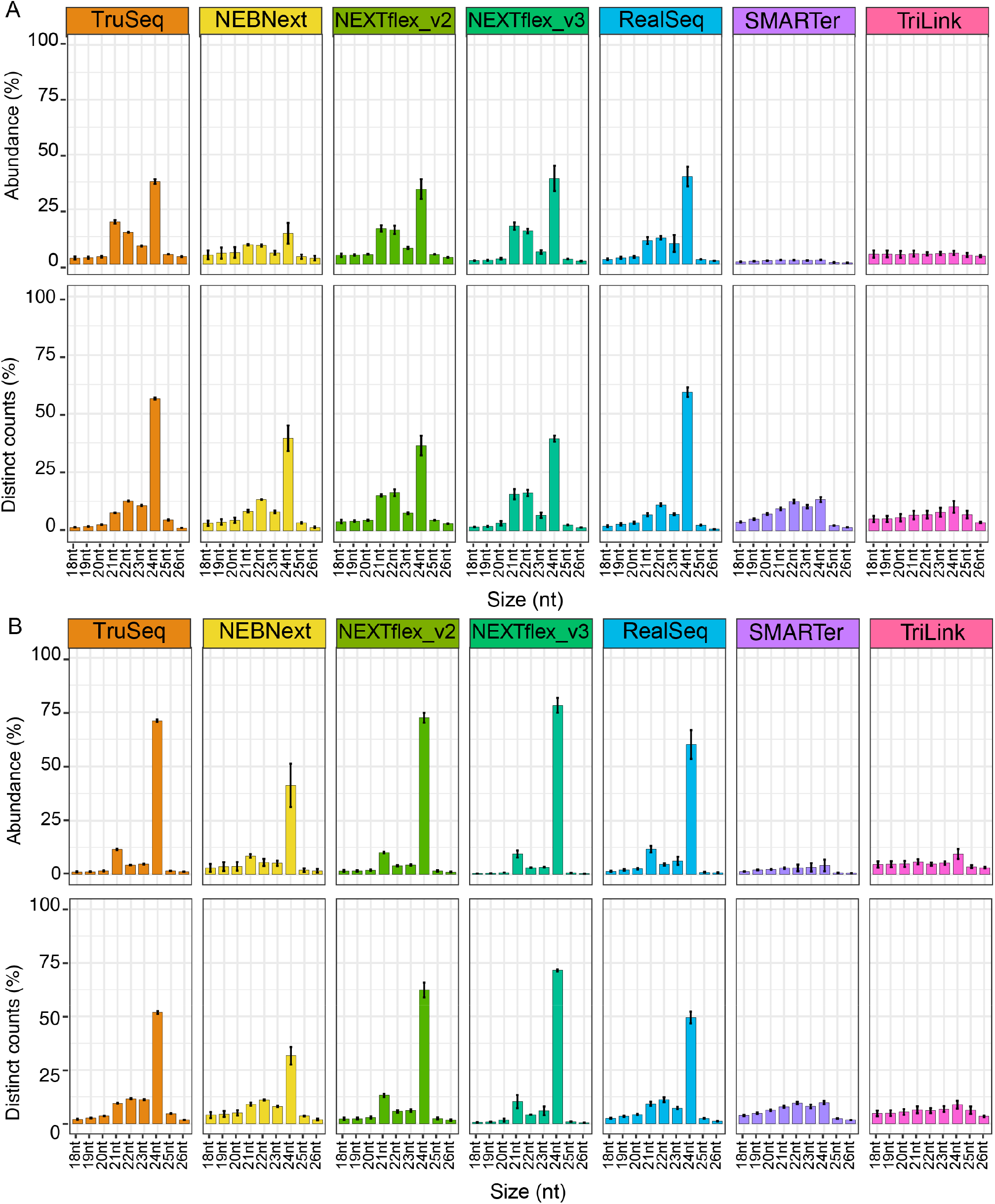
Size distribution of small RNAs mapping to the maize genome. For each plot, the x axis indicates the sRNA size (in nucleotides) and the y axis indicates the abundance (in percentage of the total number of reads), for the upper panel and distinct counts (in percentage of the total number of reads) for the lower panel. A Pre meiotic anthers, B Meiotic anthers.

We observed a few differences when looking at the abundance of the different types of sRNAs present in each library method. SMARTer and TriLink methods present a higher percentage of structural RNAs (tRNAs, rRNAs and nuclear RNAs), and a big decrease in reads that map to *PHAS* loci. TruSeq presents a higher number of reads mapping to miRNAs, and TruSeq, NEBNext, NEXTFlex and RealSeq are similar in terms of the number of reads mapping to *PHAS* loci.

### Library complexity depends on the method and starting amounts

To evaluate each library method, we generated sRNA libraries using different amounts of input RNA, ranging from 1 ng to 1000 ng. We then analyzed the number of miRNAs and *PHAS* loci identified by each library method and input, as a measure of library complexity (Figure 3). For this study, we focused our analysis on the 325 miRNAs included in the last release of miRbase (17) and the 463 21-nt and 176 24-nt *PHAS* loci identified in Zhai et al. (2015), and considered that 1 read was sufficient for an sRNA to be considered present. We observed that the number of *PHAS* loci is more consistent between the different conditions, compared to miRNAs. This is probably due to the fact that for miRNAs, we are comparing a single sRNA, and for *PHAS* loci we are comparing a set of sRNAs that map across the entire phasiRNA-producing locus.

**Figure 3:**
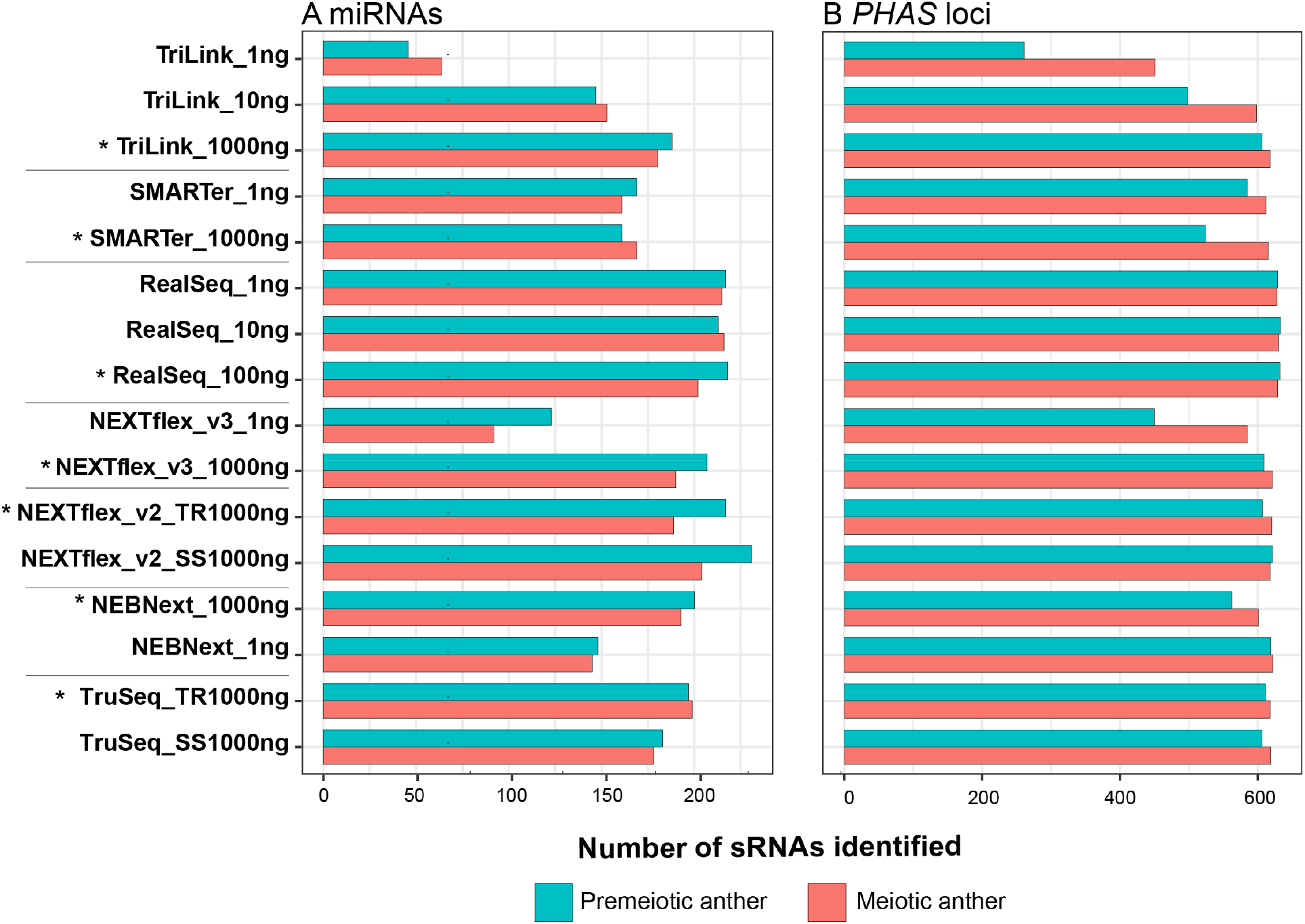
Library complexity depends on the method and starting amounts. The x axis indicates the sample, including the library preparation method and the amount of RNA used as starting material, and the y axis indicates the number of small RNAs identified. The samples labeled with a “*” used the amount recommended by the manufacturer. A miRNAs, B *PHAS* loci.

We first evaluated each commercial method using the recommended starting amount (Figure 3 – star labeled samples). We obtained the maximum complexity using NEXTFlex v2 with 1000 ng of starting material and performing size selection, followed by RealSeq with 100 ng, and NEXTFlex v3 with 1000 ng. However, as one of the aims of this study was to identify reliable methods for low input libraries, we considered low input libraries of 10 ng or 1 ng, and for both PMA and MA tissues, we obtained the maximum complexity using the RealSeq method. It is worth to note that the vast majority of miRNAs were identified in all the different construction methods, regardless of the initial RNA input amount. The results of overlapping miRNAs identified in each method with different RNA input quantities are displayed in Supplemental Figure 2.

We also wanted to assess whether different starting amounts would capture different miRNAs. We analyzed the percentage of identified miRNAs that overlap when comparing the different conditions of each method (Supplemental Figure 3). The RealSeq method had the highest overlap between recommended input and low input samples, reaching 80% or more in both tissues. Other methods, like NEBNext, were not far from this overlap percentage, reaching 65% and 75%, respectively, in meiotic and premeiotic anthers. However, we observed lower than expected overlap between low and high input samples for NEXTFlex (~50%), SMARTer (~63%) and TriLink (~26%).

We conclude that if the starting amount of material is not limiting, NEXTFlex or RealSeq methods provided adequate complexity. However, when working with low amounts of RNA, RealSeq was the only method that preserved high levels of complexity.

### Adapter ligation bias occurs at both ends of the miRNAs

To assess the ligation bias of each kit, we analyzed the nucleotide composition for each position of the miRNAs identified by each method, using samples prepared with the RNA input amount recommended by the manufacturer (Figure 4). We focused on the nucleotide composition of the first and last nucleotides, since this will determine the composition at all other positions. We observed that even if most of the identified miRNAs in each kit contained a U/T in their 5’ end, differences still exist between kits. The two extremes are represented by the TruSeq method, that identifies almost exclusively miRNAs with a 5’ U/T, and TriLink, that yields a higher diversity, including miRNAs with a U/T, G, or A as a 5’ end. Concerning the 3’ end, we observed a higher diversity than in the 5’ end. While the TruSeq method mostly identifies miRNAs starting with a C, NEXTFlex kits mostly identify miRNAs with a G at the 3’ end. In most of the methods, miRNAs with an A at the 3’ end are almost completely absent. We repeated the same analysis using only the samples prepared using 1 ng of input RNA and obtained the exact same results. These observed differences might be one of the principal reasons for the quantitative differences observed between kits.

**Figure 4:**
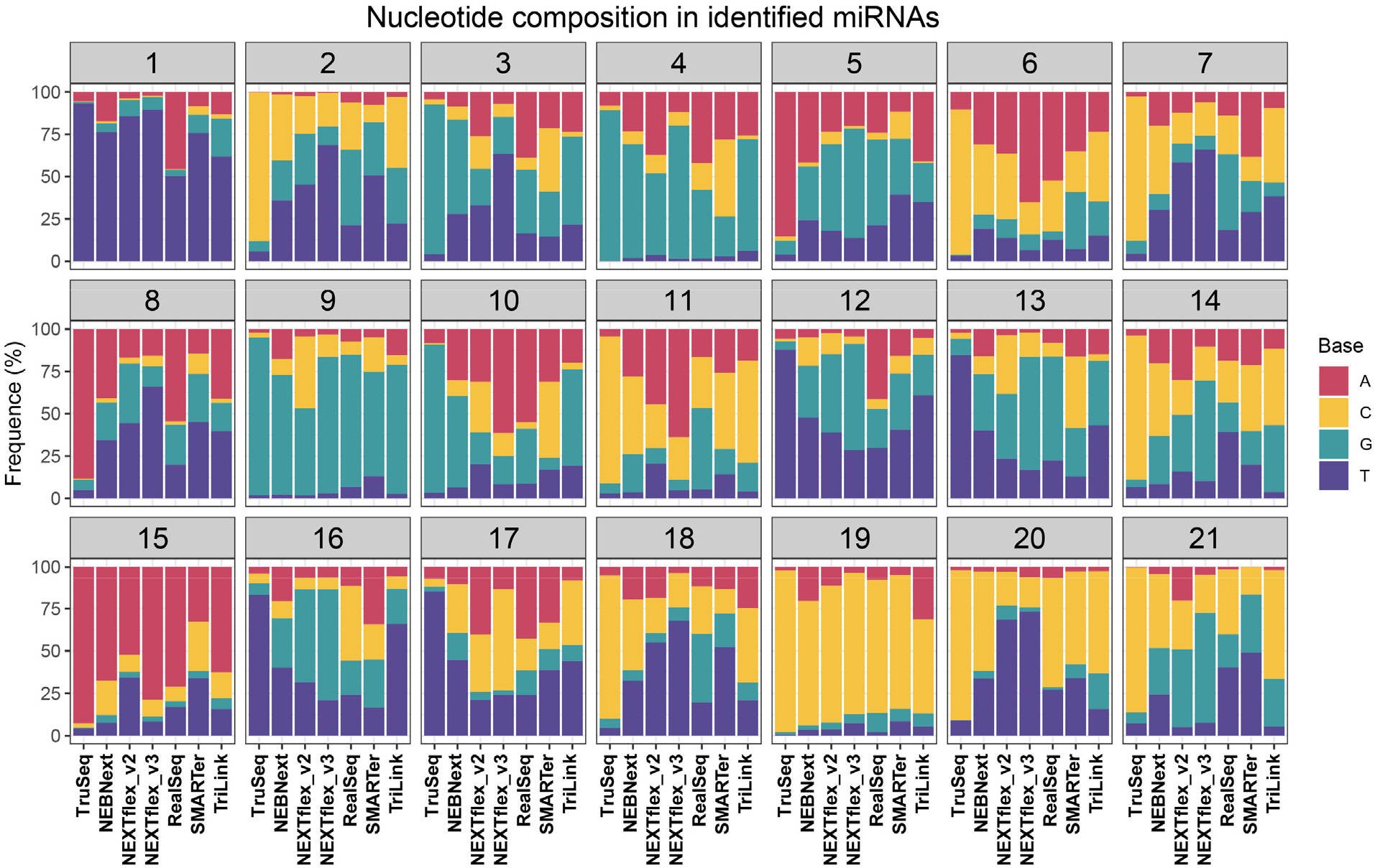
Adapter ligation bias occurs at both ends of the miRNAs. Each bar plot represents one position of the miRNA. For each plot, the x-axis indicates the library construction method, and the y-axis indicates the frequency of each nucleotide, in percentage. The data here represented includes the average of three technical replicates, using only the manufacturer’s recommended starting amount. Composition was plotted using FastQC output.

### PhasiRNA accumulation is extremely variable at all levels

PhasiRNAs are a heterogeneous group of sRNAs, either 21- or 24-nt long, generated from double-stranded RNAs in sequential phases or cycles (27). Due to their high variability of 5’ and 3’ nucleotides, phasiRNAs are less subject to bias in adapter ligation. To evaluate the ability of each method to capture phasiRNAs, we analyzed their accumulation using samples prepared with the RNA input amount recommended by the manufacturer. We then selected the most abundant 21- and 24-nt producing *PHAS* loci and studied the accumulation of each phasiRNA (labeled as “cycle” in Figure 5). We observed that phasiRNA accumulation is extremely variable when comparing between kits (Supplemental Figure 4, Supplemental table 2). As described before (28, 29), most phasiRNAs from a single *PHAS* locus accumulate to diverse levels (Supplemental Figure 4, Supplemental table 2). The biogenesis of phasiRNAs by processive activity of Dicer should yield phasiRNA duplexes from a given locus in stoichiometrically equal levels; yet, of tens of thousands of reproductive phasiRNAs only a small proportion are abundant and observed in nonstoichiometric abundances within a PHAS locus (28). This variation may be attributed to stabilization by loading in AGO proteins or interactions with targets. We observed that the abundance of each phasiRNA is largely dependent on the method used for library construction. For example, in Figure 5, we represent the abundance of two 21- and two 24-nt *PHAS* loci. We observed that the most abundant phasiRNA for each of these *PHAS* loci varies for each method. For the two 21-nt phasiRNAs, the most abundant cycles are 8 and 7, 1 and 2, 10 and 3, and 6 and 19, for NEBNext, NEXTFlex, RealSeq and TruSeq respectively. For the 24-nt *PHAS* loci, we observed similar variation in the accumulation patterns; in this case, the most abundant phasiRNAs were in positions 10 and 5, 17 and 5, 16 and 8, and 17 and 12, for NEBNext, NEXTFlex, RealSeq and TruSeq respectively. It is worth noting that the two other methods evaluated in this study, TriLink and SMARter, barely detected any phasiRNAs.

**Figure 5:**
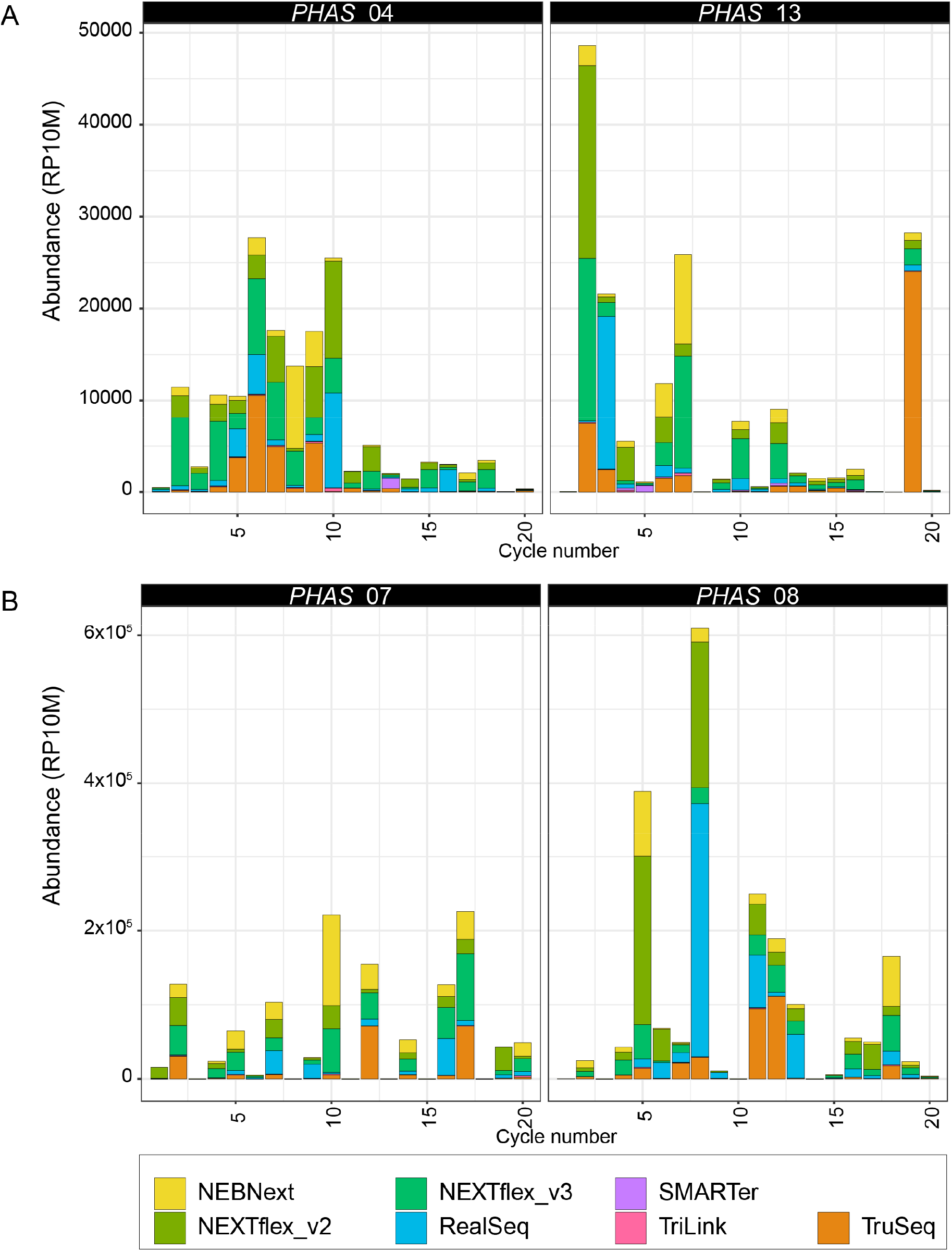
PhasiRNA accumulation is extremely variable at the levels of the *PHAS* locus, phasiRNAs, and kits. For each plot, the x-axis indicates one cycle number (corresponding to each mature phasiRNA), and the y-axis corresponds to the accumulative abundance in all kits, in reads per ten million (RP10M). For more details, see Supplemental Figure 4.

To assess whether this variability is influenced by bias in adapter ligation, we analyzed the nucleotide composition of phasiRNAs captured by each method, assessing the proportion of each nucleotide at each position (Figure 6, Supplemental Figure 5). We observed that the majority of methods capture 21-nt phasiRNAs with a cytidine (C) and 24-nt phasiRNAs with an adenosine (A) in their 5’ end (Figure 6 A). However, using the RealSeq method, more than 60% and 80% of the captured 21-nt and 24-nt phasiRNAs, respectively, had an adenosine (A) in their 5’ end. We also observed a second bias in position 19 for 21-nt phasiRNAs (Figure 6A). As described previously (30), the 19^th^ position presents a depletion of adenosine (A), that is independent of the method used for library construction. However, this depletion was not observed in any position in 24-nt phasiRNAs (Figure 6B, Supplemental Figure 5B). Additionally, the SMARTer method displayed a strong depletion of adenosine (A) at the 3’ end on both 21- and 24-nt phasiRNAs. We hypothesize that this 3’ end depletion might be due to the polyadenylation process used during library construction. These observed technical biases are consistent between replicates, and independent of the RNA amount used as input.

**Figure 6:**
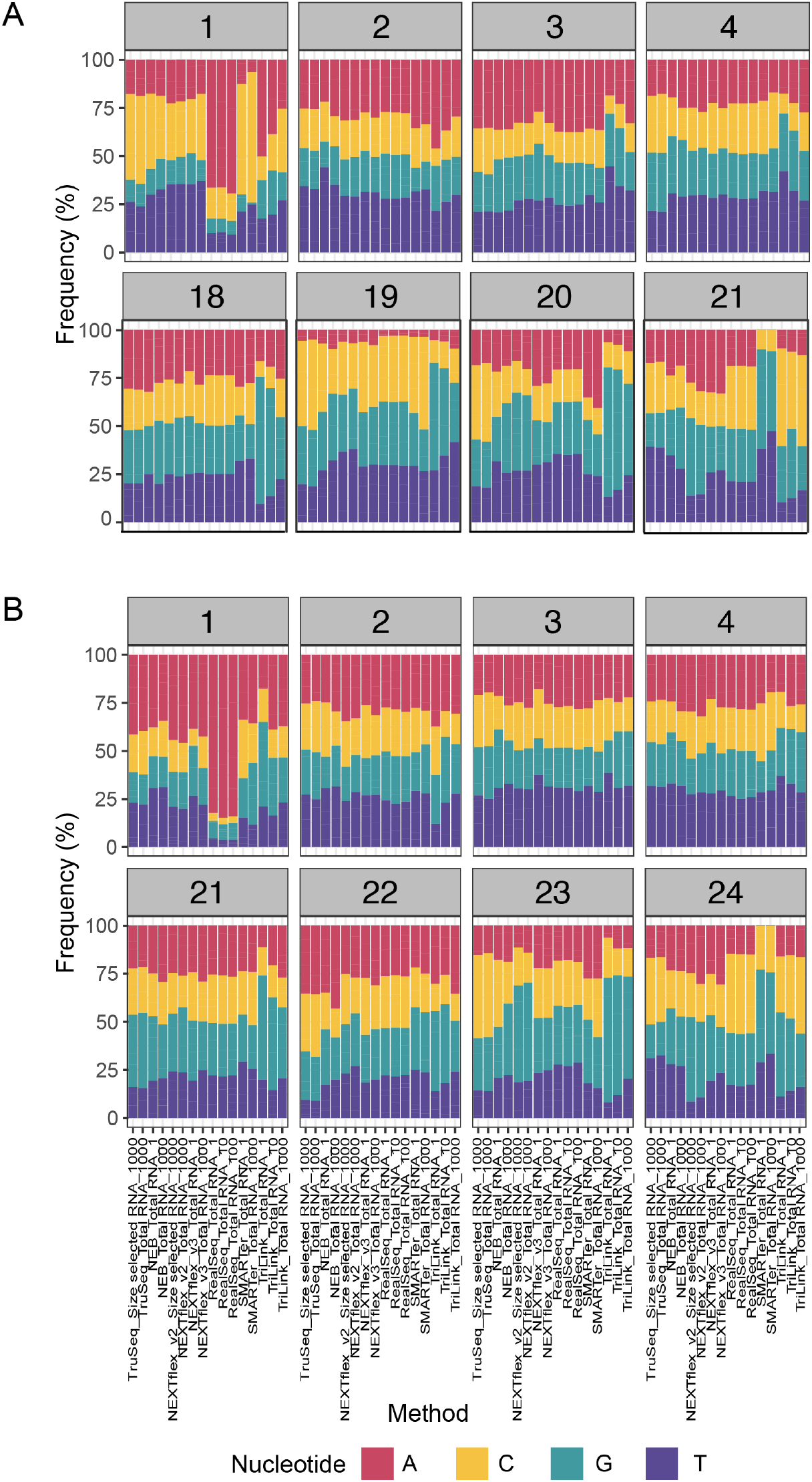
PhasiRNA accumulation is extremely variable at the levels of the *PHAS* locus, phasiRNAs, and kits. Each bar plot represents one position of the 21- (A) and 24-nt phasiRNAs (B), respectively. For each plot, the x-axis indicates the library construction method, and the y-axis indicates the frequency of each nucleotide, in percentage. The data here represented includes the average of premeiotic and meiotic anthers, with three technical replicates each. Composition was plotted using FastQC output. For simplification, only the four first and last positions are represented. For more details, see Supplemental Figure 5.

### RealSeq, TruSeq and NEXTFlex demonstrate the most similar expression profiles

To assess the correlation in the abundance of miRNAs and phasiRNAs, we used the Spearman’s rank correlation coefficient to compare the relative position of each observation within the variable. When examining sRNA abundances, a correlation of 100 indicates that the same sRNAs are ranked in the exact same positions when comparing two different methods. We calculated the correlation matrices for both types of samples, premeiotic anthers (Figure 7, upper plots) and meiotic anthers (Figure 7, lower plots), and for miRNAs (Figure 7A) and *PHAS* loci (Figure 7B). When comparing different samples within each method, most of the kits had a high correlation coefficient. This indicates that all methods of library preparation yielded highly reproducible results for mature miRNAs and *PHAS* loci summed abundances, even if the individual phasiRNA abundances are highly variable across *PHAS* loci. However, regardless of the type of tissue or the type of sRNA analyzed, RealSeq had the highest correlation coefficient, between 94% and 100%. All RealSeq samples also had a high correlation coefficient with all other methods when using samples prepared with the RNA input amount recommended by the manufacturer, indicated in the figure with an asterisk, especially with NEXTFlex and TruSeq methods.

**Figure 7:**
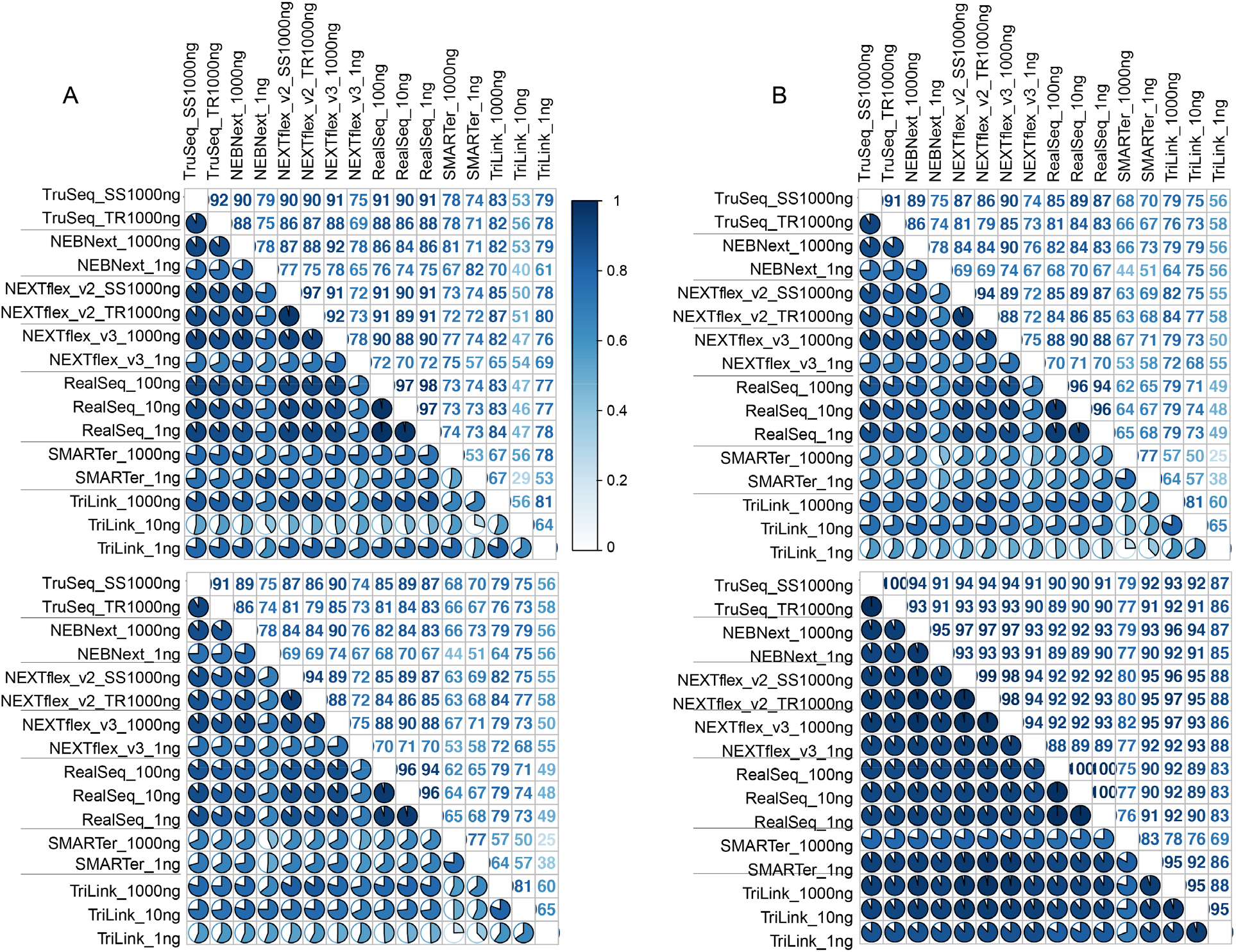
The correlation of small RNA abundances is variable across library construction methods. For each plot, the upper and lower halves (above and below the diagonal) indicate the percentage of correlation in numeric values and pie charts, respectively. The upper panels correspond to premeiotic anthers (PMA) and the bottom panels correspond to meiotic anthers (MA). The name of each sample indicates the method used for library preparation and the amount of RNA used as starting material. A. microRNAs and B. *PHAS* loci.

Another way that we assessed the impact of the library preparation methods on expression profiles was to analyze the differentially accumulated (DA) miRNAs between premeiotic and meiotic anthers. We used the DeSeq2 package to calculate the DA of miRNAs using samples prepared with the RNA input amount recommended by the manufacturer. We plotted the results as a heatmap (Figure 8) and we observed that most of the methods produce a similar DA pattern, including one block of miRNAs down-regulated (I) and one block up-regulated (II), in meiotic versus premeiotic anthers. We also observed that the clustering analysis grouped RealSeq, NEXTFlex v2 and TruSeq as the most similar methods, confirming the results obtained using correlation plots.

**Figure 8:**
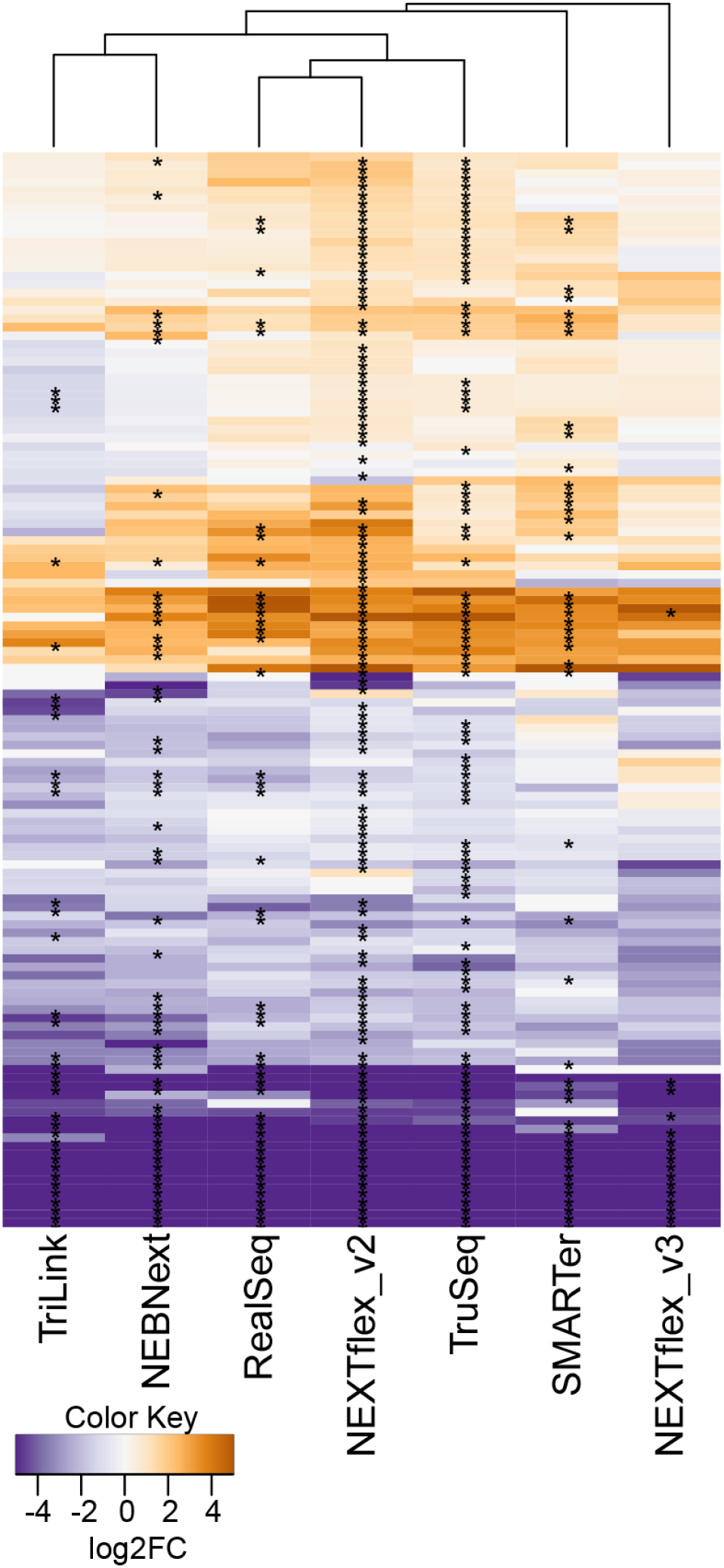
Differentially accumulated miRNAs of PMA compared to MA were consistent across most methods. The heatmap included differentially accumulated miRNAs for each construction method, comparing premeiotic anthers (PMA) to meiotic anthers (MA). The data represented here includes the average of three technical replicates, using only the manufacturer’s recommended starting amount. Statistically significant comparisons (p-value < 0,05) are indicated with a “*”.

### Identified tRFs are variable in size and origin between methods

In the last few years, the focus of sRNAs has moved from mainly miRNAs to all kinds of sRNAs. tRNA-derived fragments (tRFs) play essential roles in genome protection and transposon movement (31, 32), although the mechanism by which tRFs function is not fully known. For the purpose of this study, we considered tRFs from 19- to 24-nt, that map to tRNAs. The various library construction methods we studied revealed variable tRF accumulation patterns (Figure 9). TriLink detected the highest number of tRFs with a high majority of reads as 23 nt long, and derived from alanine (Ala) coding tRNAs. RealSeq and NEXTFlex v3 also detected high amounts of tRFs, mostly derived from leucine (LEU) and glutamic acid (GLU) tRNAs. The size of the identified tRFs was also extremely variable. RealSeq, TruSeq and NEBNext identified mostly 24-nt long tRFs while NEXTFlex 2 and TriLink identified mainly 23 nt tRFs. This extremely high variability of detected tRFs could contribute to why their function and mode of action remains unknown.

**Figure 9:**
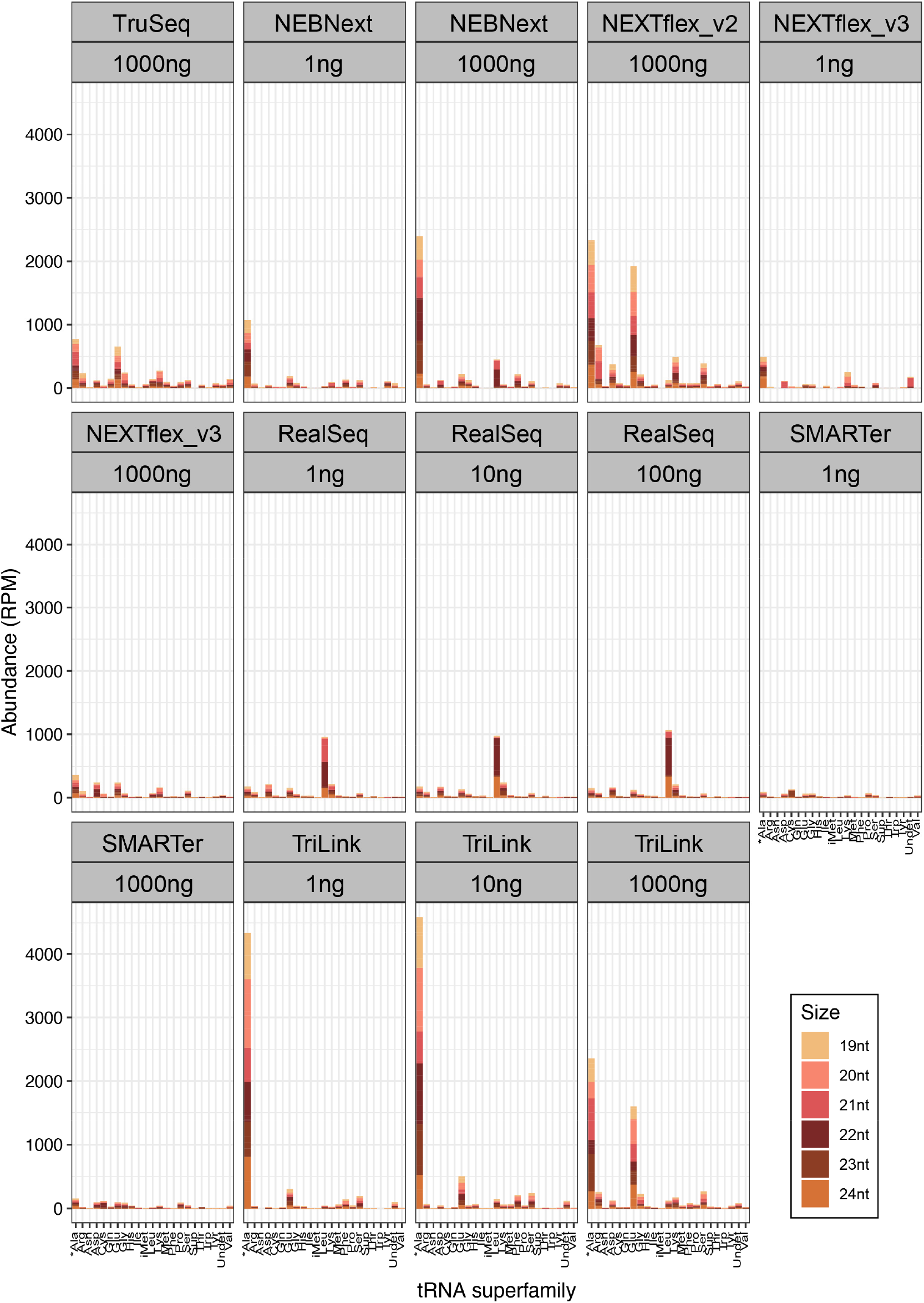
Identified tRFs are variable in size and origin between methods. Bar plot showing the size and origin of identified tRNA-derived fragments (tRFs). The x axis represents the amino acid of origin of each tRF, and the y axis represents the abundance in reads per million (RPM). The asterisk (x axis) indicates that for scaling purposes, alanine-derived tRFs are represented in reads per 10 million instead of reads per million. We also included information of the different sizes of each tRFs (from 19- to 24-nt) in different colors, as indicated in the color key.

### The size and origin of hc-siRNAs is consistent between methods

Heterochromatic siRNAs (hc-siRNAs) play an important role in transposable element silencing, stress responses and genome stability via transcriptional regulation of gene expression by RNA-directed DNA methylation (32). For the purpose of this study, we considered hc-siRNAs to be any sRNA from 21- to 24-nt long that maps to transposable elements. Due to the repetitive nature of hc-siRNAs, it is impossible to know the exact origin of each read; for that reason, we decided to group the hc-siRNAs by TE superfamilies.

We observed that all library construction methods had a similar accumulation pattern for TE-derived sRNAs, and that most of the hc-siRNAs originated from Copia, Gypsy or other LTR retrotransposons and were 24-nt in length (Figure 10). However, we observed differences in the abundance of hc-siRNAs and found that RealSeq, TruSeq and NEXTFlex methods detected more hc-siRNAs reads per million (RPM). In contrast, SMARTer and TriLink methods were only able to detect 20-30% of the hc-siRNAs reads compared to other methods (Figure 10).

**Figure 10:**
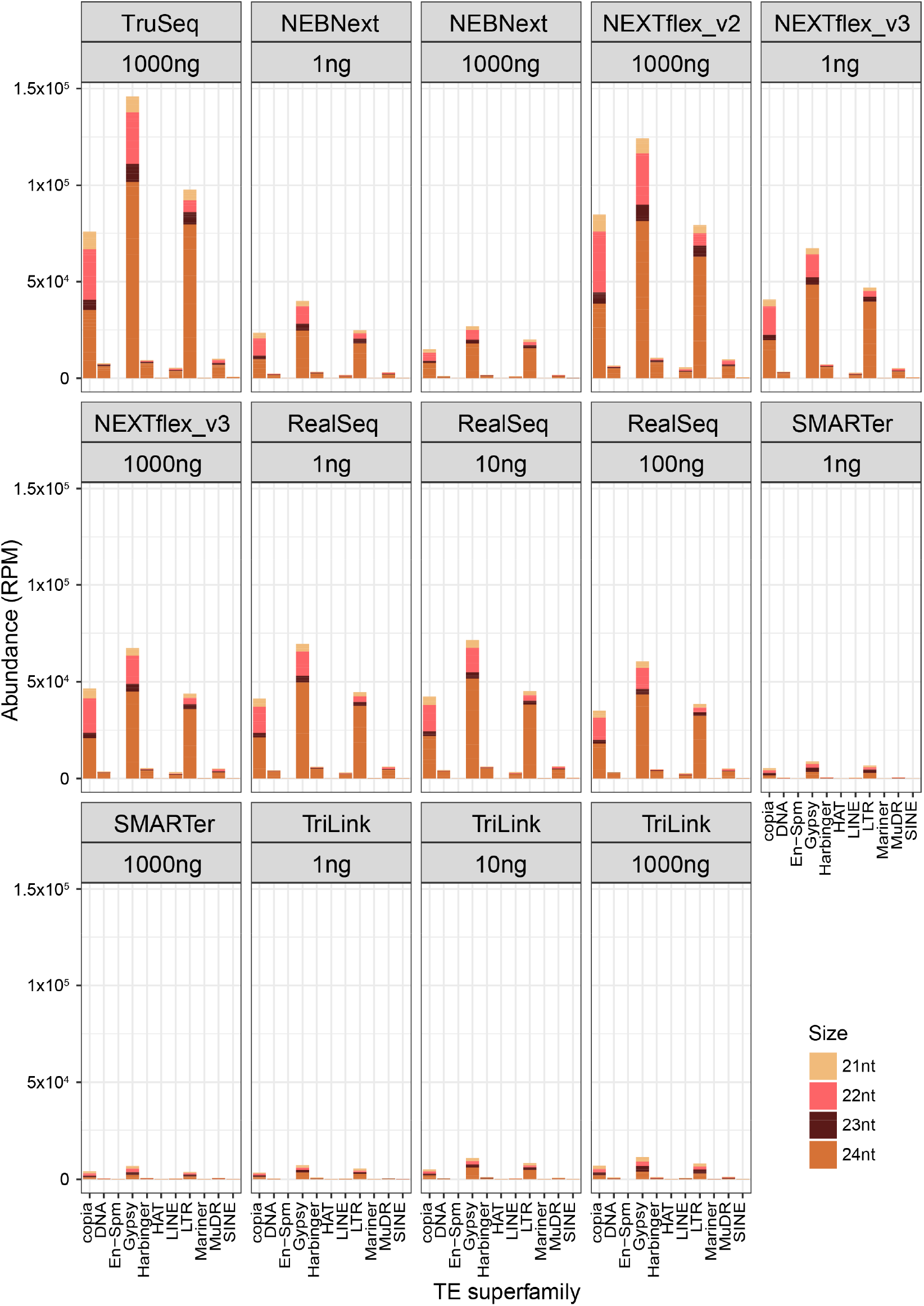
The size and origin of hcsiRNAs is consistent, independently of the method. Bar plot showing the size and origin of identified heterochromatic siRNAs (hc-siRNAs). The x axis represents the transposable element (TE) of origin of each hc-siRNA, and the y axis represents the abundance in reads per million (RPM). We also included information on the size of each hc-siRNA (from 21- to 24-nt) in different colors, indicated in the color key.

## DISCUSSION

It is known from both our work and prior studies (2–6) that several biases occur during the construction of sRNA libraries. In this study, we compared the accumulation of four different types of sRNAs in plant material using seven commercially available library construction methods. We demonstrated that ligation bias exists when using plant samples, independent of the starting RNA input amount. This is primarily caused by differences in adapter ligation efficiency. The two major factors playing a role in ligation efficiency are the secondary structure of the RNA, and the specific end nucleotides present in each RNA molecule.

We also observed strong differences in the nucleotide composition of the 5’ end of phasiRNAs, mainly for 21-nt phasiRNAs, depending on the library construction method. This type of bias has been observed previously in sequencing data (30, 33), and attributed to a preference of an Argonaute – that is, a relevant biological bias. However, our results suggest that this bias might also be technical in nature, influenced by the library construction method. Thus, the previously-observed nonstoichiometric abundances measured across several thousand of maize and rice *PHAS* loci and tens of thousands reproductive phasiRNAs (28) likely reflects a combination of biological and technical factors.

We conclude that each library construction method might be more or less adequate, depending on the RNA of interest, but that some methods work better for specific sRNAs. NEXTFlex, RealSeq and TruSeq are able to detect a higher number of reads for microRNAs and phasiRNAs. However, for abundant RNAs like ribosomal RNAs and transfer RNAs, TriLink and SMARTer libraries are enriched in these categories. Alternatively, for studies focused on nucleolar RNAs, we observed a slight enrichment in these when using TriLink. We also observed smaller differences in the accumulation of hc-siRNAs when comparing construction methods; this is probably due to the repetitive and diverse nature of these sRNAs.

In recent years, many new and different methods for sRNA library construction are commercially available. Most of these new protocols aim to more accurately reflect sRNA composition by focusing on reducing one identified bias, while leaving other biases unaddressed. While preparing this manuscript, a new method based on randomized splint ligation was published that involves a double-stranded adapter with a short, singlestranded degenerate extension (34). Other strategies have been developed to ensure absolute normalization of the sRNA library. For example, it has been proposed to use single-stranded RNA 5’ monophosphate and 2’-O-methyl oligonucleotides as spike-ins for plant samples (35). In our opinion, and based on the results described here, the new approaches might be combined to provide a new library construction method that addresses all known biases, and thus provide a more reliable picture of the sRNA population in plants. This would ideally include a circular adapter, with four degenerated nucleotides at each end, while using an artificial spike-in added to the RNA before library construction.

## DATA AVAILABILITY

All raw and processed sequencing data generated in this study have been submitted to the NCBI Gene Expression Omnibus (GEO; http://www.ncbi.nlm.nih.gov/geo/).

## SUPPLEMENTARY DATA

Supplementary Data are available at NAR online.

## ACKNOWLEDGEMENTS

We thank Joanna Friesner for the critical review of the manuscript, Mayumi Nakano for assistance with data handling as well as all the members of the Meyers lab for support and helpful discussions.

---

**Supplemental Table 1.** List of the small RNA library preparation kits tested in this study.

| Company | Kit | Cat. # | Approach |
| --- | --- | --- | --- |
| Illumina | TruSeq Small RNA Library Prep | RS-200-0012 | Adapter ligation |
| Bioo Scientific | NEXTFlex™ Small RNA-seq v2 | 5132-03 | Adapter 4N-ends |
|  | NEXTFlex™ Small RNA-seq v3 | 5132-05 | Adapter 4N-ends |
| TriLink Biotechnologies | CleanTag™ Small RNA Library Prep | L-3206 | Chemically-modified adapters |
| New England BioLabs | NEBNext® Small RNA Library Prep | E7300S | Adapter ligation |
| Clontech | SMARTer smRNA-Seq | 635029 | Ligation-free |
| Somagenics | RealSeq®-AC miRNA Library Kit | 500-00048 | Circular adapter |

**Supplemental Table 2.** Genome coordinates of the most abundant 21- and 24nt *PHAS* loci

| Type | ID | Chromosome | Coordinate | Strand |
| --- | --- | --- | --- | --- |
| 21-PHAS | PHAS_01 | 2 | 141966025 | c |
| 21-PHAS | PHAS_02 | 1 | 164855798 | c |
| 21-PHAS | PHAS_03 | 2 | 55560458 | c |
| 21-PHAS | PHAS_04 | 4 | 5591406 | c |
| 21-PHAS | PHAS_05 | 4 | 3571530 | c |
| 21-PHAS | PHAS_06 | 7 | 21678443 | c |
| 21-PHAS | PHAS_07 | 7 | 151055912 | c |
| 21-PHAS | PHAS_08 | 6 | 161445681 | w |
| 21-PHAS | PHAS_09 | 5 | 42512376 | w |
| 21-PHAS | PHAS_10 | 1 | 180276272 | c |
| 21-PHAS | PHAS_11 | 1 | 289764604 | w |
| 21-PHAS | PHAS_12 | 8 | 164154608 | w |
| 21-PHAS | PHAS_13 | 9 | 22976471 | c |
| 21-PHAS | PHAS_14 | 9 | 23007048 | c |
| 21-PHAS | PHAS_15 | 1 | 288678320 | c |
| 21-PHAS | PHAS_16 | 2 | 54625972 | c |
| 21-PHAS | PHAS_17 | 2 | 141451182 | w |
| 24-PHAS | PHAS_01 | 8 | 73307216 | w |
| 24-PHAS | PHAS_02 | 4 | 14576300 | w |
| 24-PHAS | PHAS_03 | 2 | 210452582 | w |
| 24-PHAS | PHAS_04 | 4 | 7689031 | c |
| 24-PHAS | PHAS_05 | 5 | 162150273 | c |
| 24-PHAS | PHAS_06 | 1 | 185480423 | w |
| 24-PHAS | PHAS_07 | 1 | 179386787 | w |
| 24-PHAS | PHAS_08 | 1 | 179381407 | c |
| 24-PHAS | PHAS_09 | 4 | 4710308 | c |
| 24-PHAS | PHAS_10 | 5 | 217208612 | w |
| 24-PHAS | PHAS_11 | 10 | 71847137 | w |
| 24-PHAS | PHAS_12 | 4 | 4630713 | c |
| 24-PHAS | PHAS_13 | 10 | 74515334 | w |
| 24-PHAS | PHAS_14 | 6 | 31982093 | c |
| 24-PHAS | PHAS_15 | 10 | 74790963 | w |

## SUPPLEMENTAL FIGURE LEGENDS

**Supplemental Figure 1.**
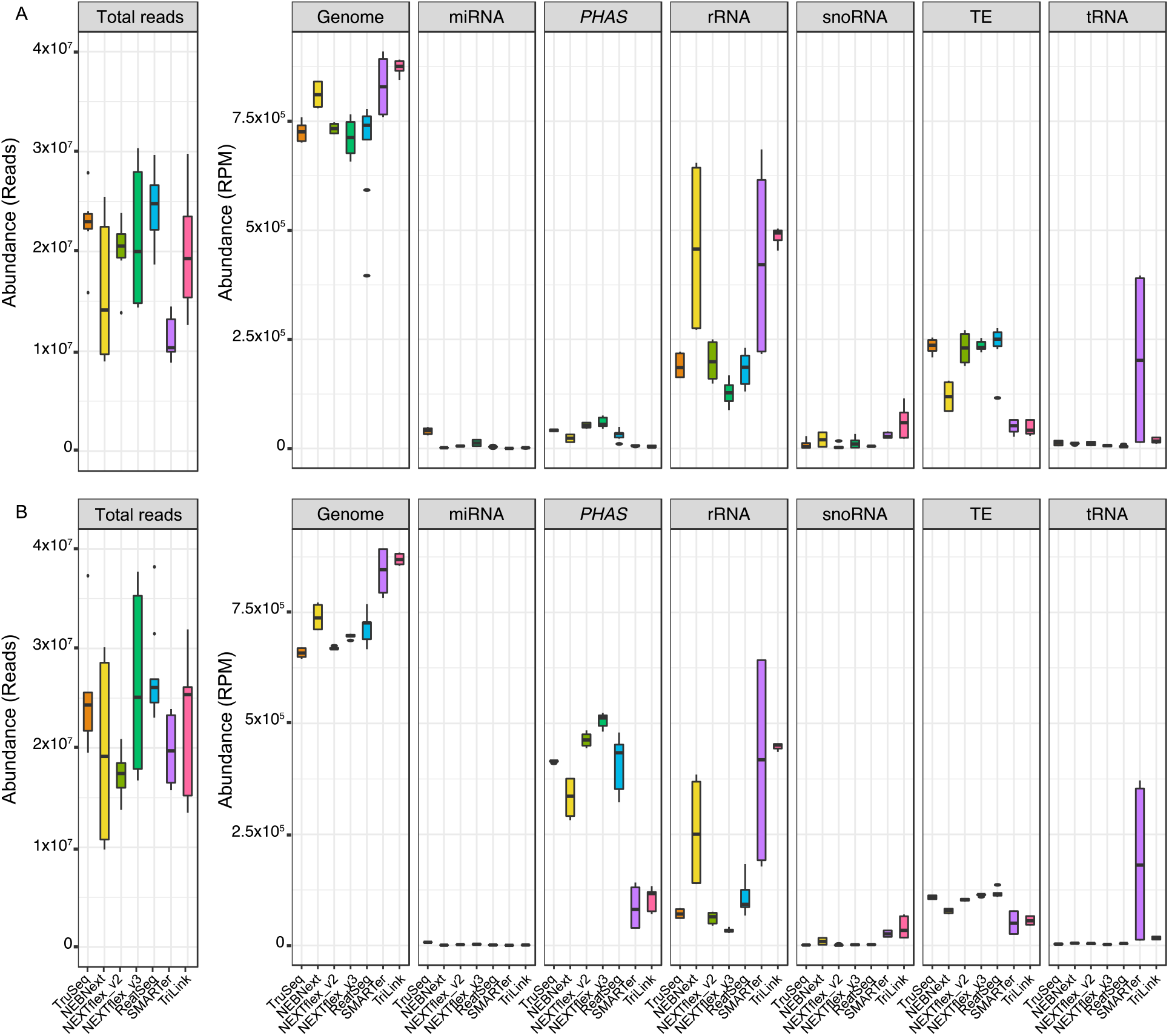
General mapping statistics of sRNA libraries. A Premeiotic anther samples. B Meiotic anther samples. The eight boxplots show the sample variation for the total number of reads, genome mapped reads (in reads per million RPM), and the abundance (in RPM) of each feature (including miRNAs, *PHAS* loci, rRNAs, snRNAs & snoRNAs, transposable elements and tRNAs) for each different library construction method. For each plot, the x-axis represents the construction methods, and the y-axis represents the abundance.

**Supplemental Figure 2:**
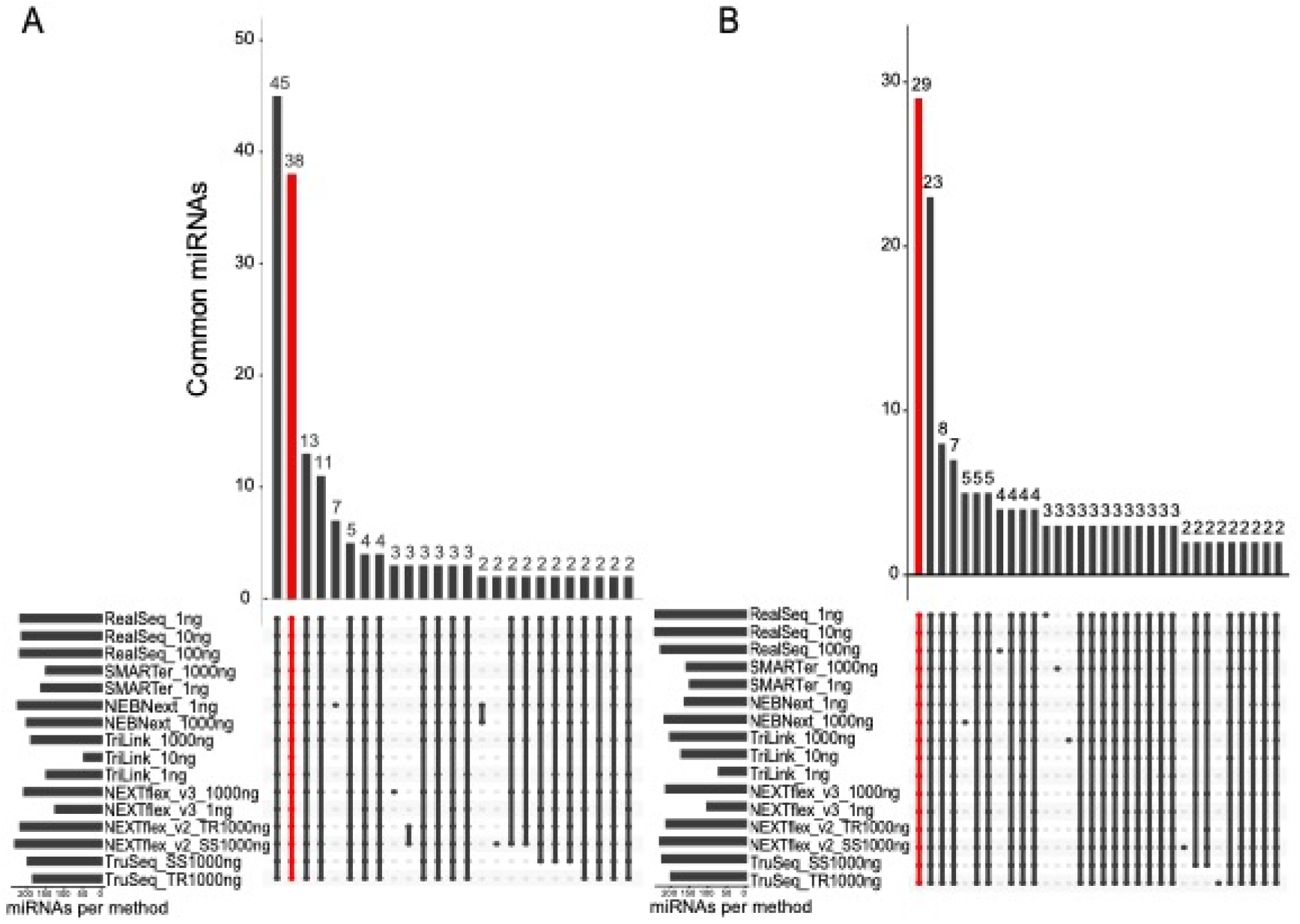
Overlap miRNA identification for each library method and RNA input amount. UpSet plots representing the overlap of identified miRNAs for each library construction and RNA input amount. Labelled in red are the miRNAs that were identified in all the different conditions. A Premeiotic anthers, B Meiotic anthers.

**Supplemental Figure 3:**
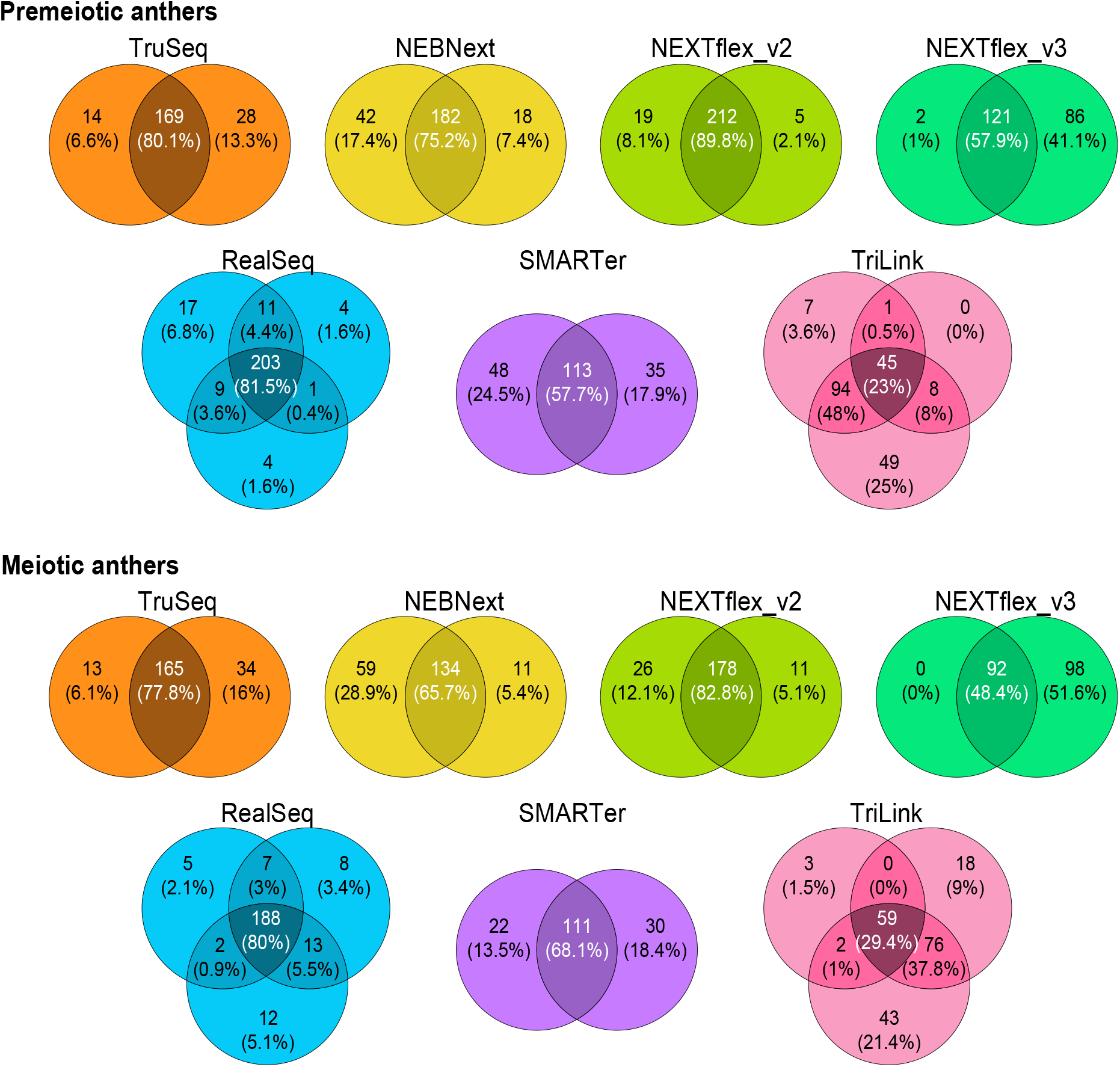
Overlap in miRNA identification for different methods, comparing different amounts of starting material. Venn diagrams representing the overlap of identified miRNAs using all the different conditions for each method.

**Supplemental Figure 4A:**
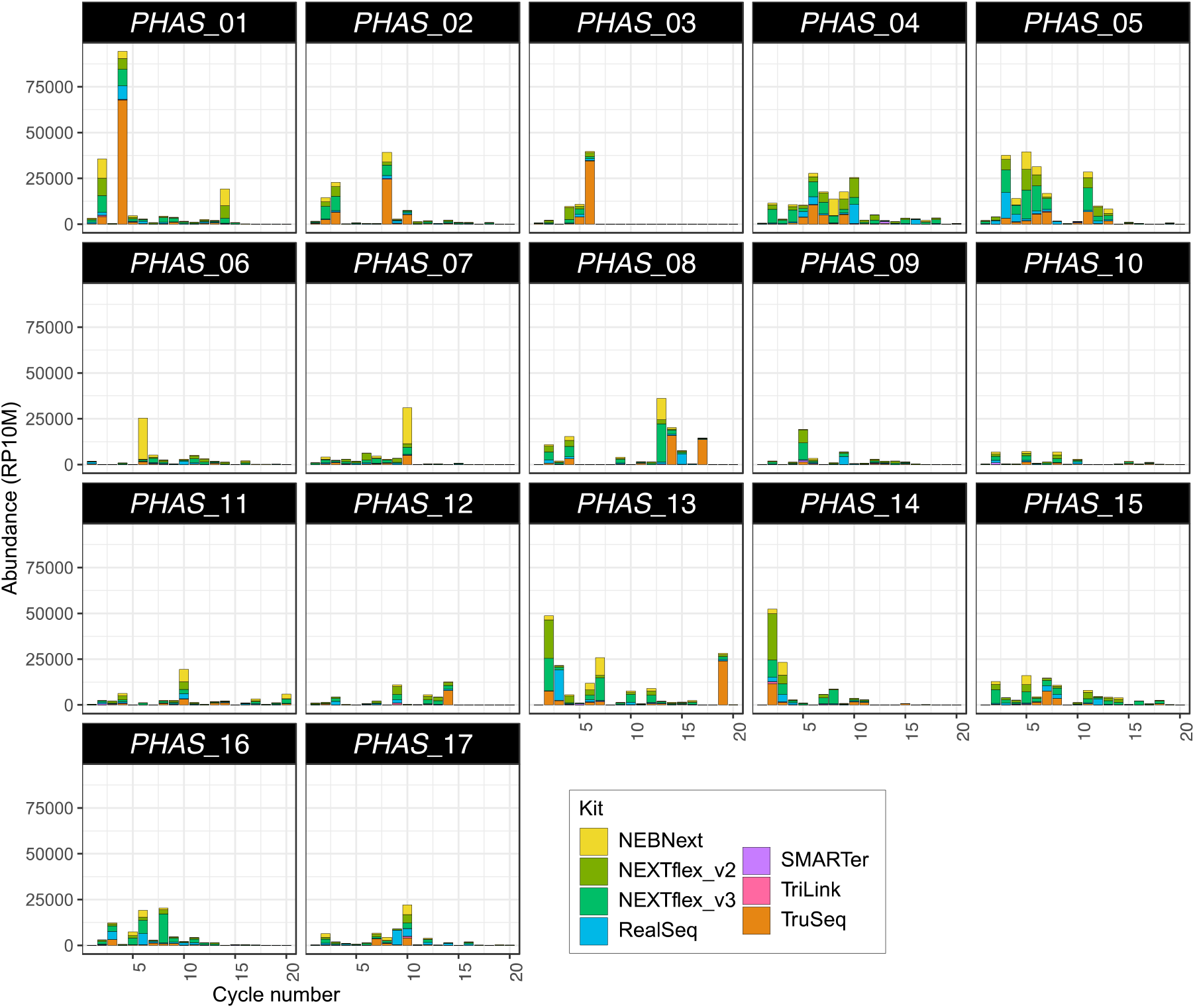
Accumulation profiles of the most abundant 21-nt phasiRNAs in each method. For each plot, the x-axis indicates one cycle number (corresponding to each mature phasiRNA), and the y-axis corresponds to the accumulative abundance in all kits, in reads per ten million (RP10M).

**Supplemental Figure 4B:**
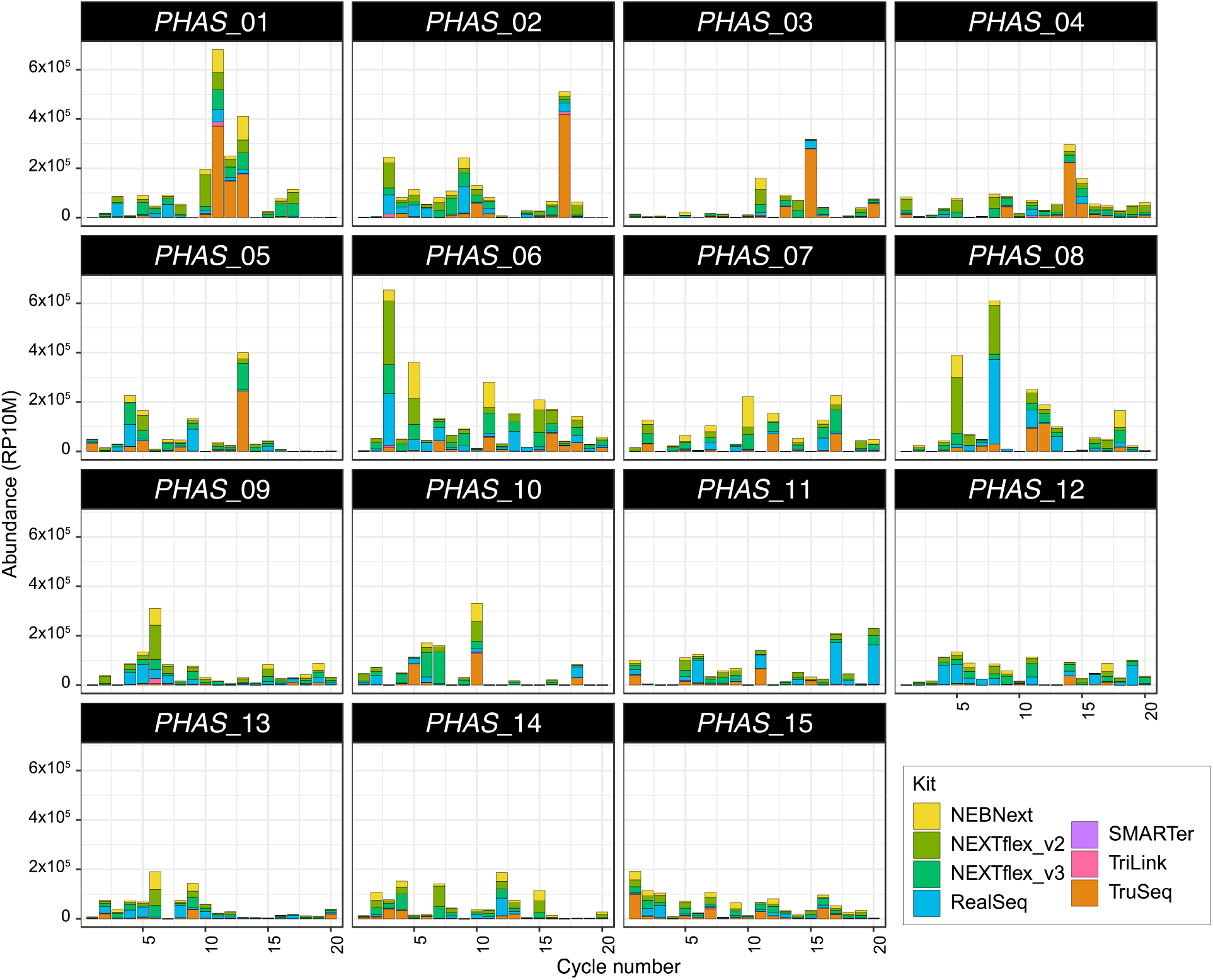
Accumulation profiles of the most abundant 24-nt phasiRNAs in each method. For each plot, the x-axis indicates one cycle number (corresponding to each mature phasiRNA), and the y-axis corresponds to the accumulative abundance in all kits, in reads per ten million (RP10M).

**Supplemental Figure 5A:**
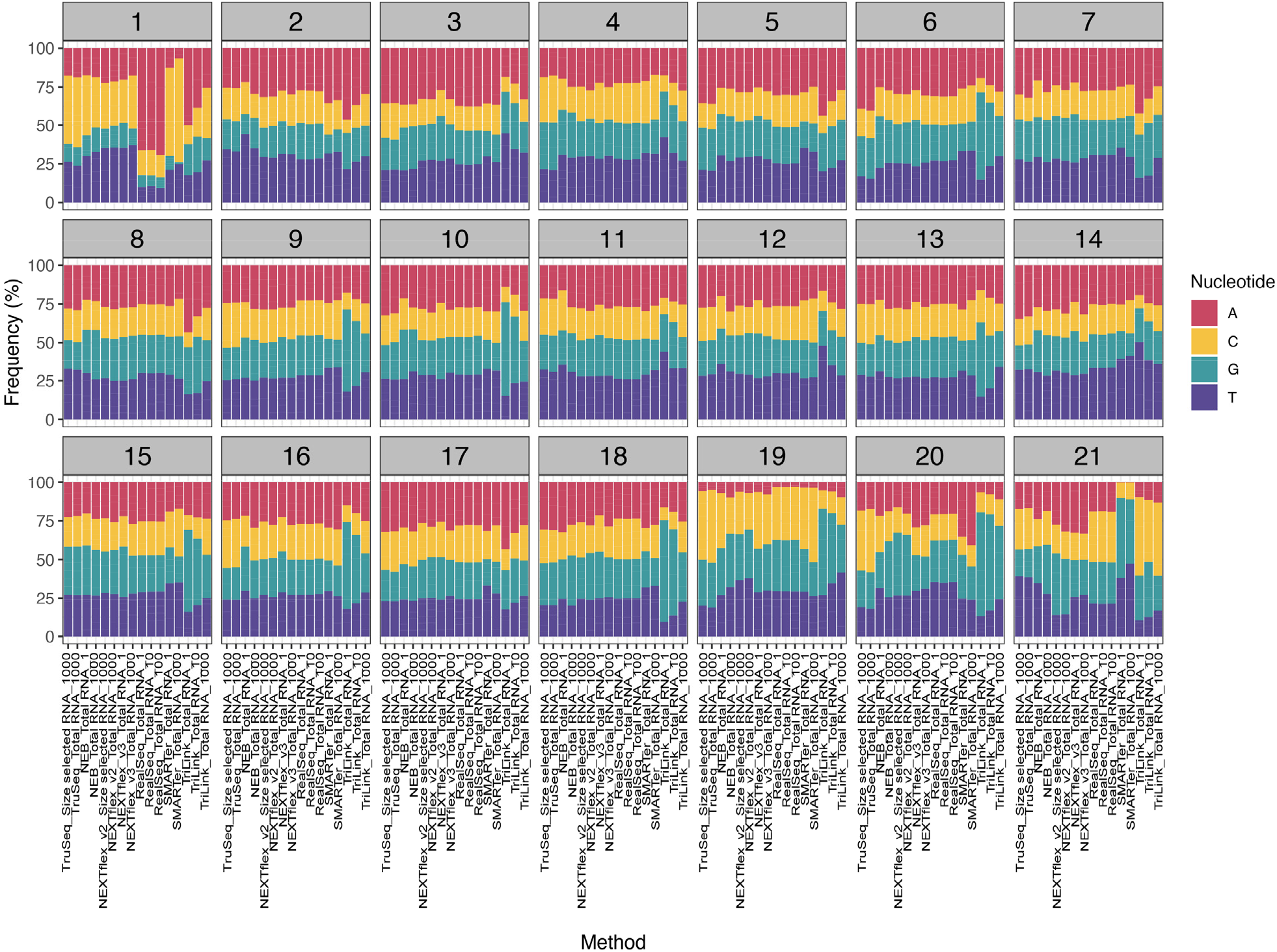
Nucleotide composition of 21-nt phasiRNAs. Each bar plot represents one position of the 21-nt phasiRNAs. For each plot, the x-axis indicates the library construction method, and the y-axis indicates the frequency of each nucleotide, in percentage. The data here represented includes the average of premeiotic and meiotic anthers, with three technical replicates each. Composition was plotted using FastQC output.

**Supplemental Figure 5B:**
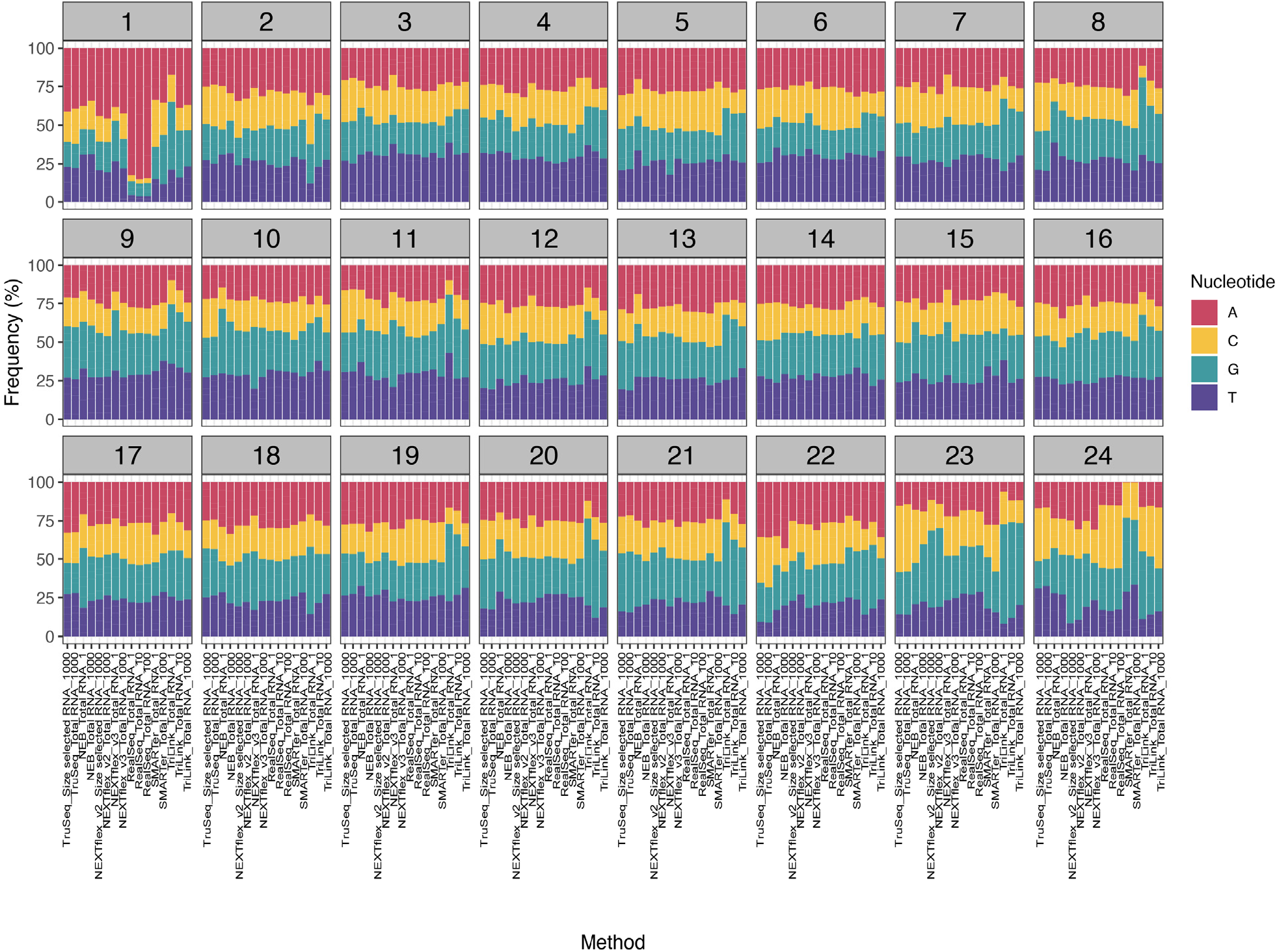
Nucleotide composition of 24-nt phasiRNAs. Each bar plot represents one position of the 24-nt phasiRNAs. For each plot, the x-axis indicates the library construction method, and the y-axis indicates the frequency of each nucleotide, in percentage. The data here represented includes the average of premeiotic and meiotic anthers, with three technical replicates each. Composition was plotted using FastQC output.

## Supplemental Methods

### Library preparation details

#### Illumina – TruSeq Small RNA Library Preparation Kit

This kit takes advantage of the natural structure of miRNAs, which have a 5’-phosphate and a 3’-hydroxyl group originated after processing of the precursor miRNAs. In summary, the protocol is based on the ligation of adapters to the 3’ and 5’ ends, respectively, followed by the synthesis of the first strand and PCR amplification with addition of barcodes allowing for multiplexing. For this kit, two amounts of input RNA were used: 1 μg of total RNA was used directly, and 2 μg of total RNA were used for 20 to 30 nt small RNA size selection on a 15% Urea TBE Polyacrylamide gel (Supplemental Table 1) as described previously (14). Thirteen PCR cycles were used for both RNA input amounts. The final cDNA libraries were size selected on 6% PAGE gels.

#### Somagenics – RealSeq™ miRNA Library Kit

This kit uses a single adapter-based approach with subsequent circularization, to reduce incorporation bias. Three amounts of input RNA were used: 100 ng, 10 ng and 1 ng of total RNA was used directly. The libraries were PCR amplified with 13, 16 and 19 cycles, respectively and were size selected on 6% PAGE gels.

#### Bioo Scientific – NEXTflex™ Small RNA-seq v2 and v3 Kits

This kit uses an adapter-based approach in which adapters with randomized bases at the ligation junctions are used to reduce ligation-associated bias. For the NEXTflex™ v2 kit, two amounts of input RNA were used: 1 μg of total RNA was used directly, and 2 μg of total RNA were used for 20-30 nt sRNA size selection on a 15% Urea TBE Polyacrylamide gel (Supplemental Table 1). The libraries were PCR amplified with 18 and 12 cycles, respectively, and both were size selected on 6% PAGE gels. For the NEXTflex™ v3, two amounts of input RNA were used: 1 ng and 1 μg of total RNA were used directly for library construction. The libraries were PCR amplified with 25 and 12 cycles, respectively. The libraries constructed with the 1 ng total RNA were size selected on 6% PAGE gels, and those prepared with 1 μg were size selected using the NEXTflex™ Cleanup Beads.

#### TriLink Biotechnologies – CleanTag™ Small RNA Library Prep Kit

This kit uses chemically modified adapters for reduced formation of adapter dimers. For this kit, three amounts of input RNA were used: 1 ng, 10 ng, and 1 μg (Supplemental Table 1). The small RNA library preparation was performed following the manufacturer’s protocol. The libraries were PCR amplified with 21, 18, and 12 cycles, respectively. The final libraries were size selected using AMPure XP Purification Beads (Beckman Coulter, cat. # A63881) following the manufacturer’s protocol.

#### New England BioLabs (NEB) – NEBNext^®^ Small RNA Library Prep Kit

In this kit, the 3’ adapter ligation to small RNAs is followed by a step to anneal the primer for reverse transcription. After this step, 5’ adapter ligation occurs, followed by the reverse transcription and final PCR amplification. Two total RNA input amounts were tested with this kit, 1 ng and 1 μg. Although the lowest input amount recommended by the manufacturer is 100 ng, for comparison purposes, we tested 1 ng of total RNA as the lowest input amount for this kit. The final libraries were size selected using AMPure XP Purification Beads (Beckman Coulter, cat. # A63881) as per the manufacturer’s instructions.

#### Clontech – SMARTer smRNA-Seq Kit

This kit uses the Clontech’s proprietary SMART (switching mechanism at the 5’ end of RNA template) technology, which first polyadenylates small RNAs, and then followed by ligation-free steps, adds the oligo dT primer containing the 3’ adapter. When the specific reverse transcriptase reaches the 5’ end of each template, it adds non-template nucleotides (usually three nucleotides) which are bound by an LNA oligo (SMART smRNA Oligo). Then, when the reverse transcriptase switches the template, it uses the LNA oligo as template to add the 5’ adapter. Indexes are added in the next step when the final library is PCR amplified. The total RNA input amounts tested for this kit are 1 ng and 1 μg (Supplemental Table 1). The number of PCR cycles used for each total RNA input amount tested was 17 and 8 cycles, respectively. The final libraries were size selected using AMPure XP Purification Beads (Beckman Coulter, cat. # A63881) as per the manufacturer’s instructions.

## REFERENCES

1. Jayaprakash, A.D., Jabado, O., Brown, B.D. and Sachidanandam, R. (2011) Identification and remediation of biases in the activity of RNA ligases in small-RNA deep sequencing. Nucleic Acids Res., 39, e141.

2. Yeri, A., Courtright, A., Danielson, K., Hutchins, E., Alsop, E., Carlson, E., Hsieh, M., Ziegler, O., Das, A., Shah, R.V., et al. (2018) Evaluation of commercially available small RNASeq library preparation kits using low input RNA. BMC Genomics, 19, 331.

3. Barberán-Soler, S., Vo, J.M., Hogans, R.E., Dallas, A., Johnston, B.H. and Kazakov, S.A. (2018) Decreasing miRNA sequencing bias using a single adapter and circularization approach. Genome Biol., 19, 105.

4. Kim, H., Kim, J., Kim, K., Chang, H., You, K. and Kim, V.N. (2019) Bias-minimized quantification of microRNA reveals widespread alternative processing and 3’ end modification. Nucleic Acids Res., 47, 2630–2640.

5. Baran-Gale, J., Kurtz, C.L., Erdos, M.R., Sison, C., Young, A., Fannin, E.E., Chines, P.S. and Sethupathy, P. (2015) Addressing Bias in Small RNA Library Preparation for Sequencing: A New Protocol Recovers MicroRNAs that Evade Capture by Current Methods. Front. Genet., 6, 352.

6. Wright, C., Rajpurohit, A., Burke, E.E., Williams, C., Collado-Torres, L., Kimos, M., Brandon, N.J., Cross, A.J., Jaffe, A.E., Weinberger, D.R., et al. (2019) Comprehensive assessment of multiple biases in small RNA sequencing reveals significant differences in the performance of widely used methods. BMC Genomics, 20, 513.

7. Rogers, K. and Chen, X. (2013) Biogenesis, turnover, and mode of action of plant microRNAs. Plant Cell, 25, 2383–2399.

8. Axtell, M.J. (2013) Classification and comparison of small RNAs from plants. Annu. Rev. Plant Biol., 64, 137–159.

9. Chen, C., Zeng, Z., Liu, Z. and Xia, R. (2018) Small RNAs, emerging regulators critical for the development of horticultural traits. Horticulture Research, 5, 6–8.

10. Zhai, J., Zhang, H., Arikit, S., Huang, K., Nan, G.-L., Walbot, V. and Meyers, B.C. (2015) Spatiotemporally dynamic, cell-type–dependent premeiotic and meiotic phasiRNAs in maize anthers. Proceedings of the National Academy of Sciences, 112, 201418918.

11. Xia, R., Xu, J., Arikit, S. and Meyers, B.C. (2015) Extensive families of miRNAs and PHAS loci in Norway spruce demonstrate the origins of complex phasiRNA networks in seed plants. Mol. Biol. Evol., 32, 2905–2918.

12. Teng, C., Zhang, H., Hammond, R., Huang, K., Meyers, B.C. and Walbot, V. (2020) Dicer-like 5 deficiency confers temperature-sensitive male sterility in maize. Nature Communications, 11.

13. Song, X., Li, P., Zhai, J., Zhou, M., Ma, L., Liu, B., Jeong, D.-H., Nakano, M., Cao, S., Liu, C., et al. (2012) Roles of DCL4 and DCL3b in rice phased small RNA biogenesis. Plant J., 69, 462–474.

14. Mathioni, S.M., Kakrana, A. and Meyers, B.C. (2017) Characterization of Plant Small RNAs by Next Generation Sequencing. Curr. Protoc. Plant Biol, 2, 39–63.

15. Bolger, A.M., Lohse, M. and Usadel, B. (2014) Trimmomatic: A flexible trimmer for Illumina sequence data. Bioinformatics, 30, 2114–2120.

16. Jiao, Y., Peluso, P., Shi, J., Liang, T., Stitzer, M.C., Wang, B., Campbell, M.S., Stein, J.C., Wei, X., Chin, C.-S., et al. (2017) Improved maize reference genome with single-molecule technologies. Nature, 546, 524–527.

17. Kozomara, A. and Griffiths-Jones, S. (2014) miRBase: Annotating high confidence microRNAs using deep sequencing data. Nucleic Acids Res., 42, 68–73.

18. Langmead, B. and Salzberg, S.L. (2013) Langmead. 2013. Bowtie2. Nat. Methods, 9, 357–359.

19. Love, M.I., Huber, W. and Anders, S. (2014) Moderated estimation of fold change and dispersion for RNA-seq data with DESeq2. Genome Biol., 15, 1–21.

20. RStudio Team (2020) RStudio: Integrated Development for R. RStudio.

21. Wickham, H. (2009) ggplot2: Elegant Graphics for Data Analysis Springer-Verlag New York.

22. Auguie, B. and Antonov, A. (2017) gridExtra: miscellaneous functions for ‘grid’ graphics. R package version, 2, 602.

23. Wei, T. and Simko, V. (2016) The corrplot package. R Core Team.

24. Lex, A., Gehlenborg, N., Strobelt, H., Vuillemot, R. and Pfister, H. (2014) UpSet: Visualization of Intersecting Sets. IEEE Trans. Vis. Comput. Graph., 20, 1983–1992.

25. Portwood, J.L., 2nd, Woodhouse, M.R., Cannon, E.K., Gardiner, J.M., Harper, L.C., Schaeffer, M.L., Walsh, J.R., Sen, T.Z., Cho, K.T., Schott, D.A., et al. (2019) MaizeGDB 2018: the maize multi-genome genetics and genomics database. Nucleic Acids Res., 47, D1146–D1154.

26. Nobuta, K., Lu, C., Shrivastava, R., Pillay, M., De Paoli, E., Accerbi, M., Arteaga-Vazquez, M., Sidorenko, L., Jeong, D.-H., Yen, Y., et al. (2008) Distinct size distribution of endogenous siRNAs in maize: Evidence from deep sequencing in the mop1-1 mutant. Proc. Natl. Acad. Sci. U. S. A., 105, 14958–14963.

27. Xia, R., Chen, C., Pokhrel, S., Ma, W., Huang, K., Patel, P., Wang, F., Xu, J., Liu, Z., Li, J., et al. (2019) 24-nt reproductive phasiRNAs are broadly present in angiosperms. Nat. Commun., 10, 627.

28. Tamim, S., Cai, Z., Mathioni, S.M., Zhai, J., Teng, C., Zhang, Q. and Meyers, B.C. (2018) Cis-directed cleavage and nonstoichiometric abundances of 21-nucleotide reproductive phased small interfering RNAs in grasses. New Phytol., 220, 865–877.

29. López-Dolz, L., Spada, M., Daròs, J.-A. and Carbonell, A. (2020) Fine-tune control of targeted RNAi efficacy by plant artificial small RNAs. Nucleic Acids Res., 10.1093/nar/gkaa343.

30. Patel, P., Mathioni, S., Kakrana, A., Shatkay, H. and Meyers, B.C. (2018) Reproductive phasiRNAs in grasses are compositionally distinct from other classes of small RNAs. New Phytol., 220, 851–864.

31. Martinez, G., Choudury, S.G. and Slotkin, R.K. (2017) TRNA-derived small RNAs target transposable element transcripts. Nucleic Acids Res., 45, 5142–5152.

32. Sigman, M.J. and Slotkin, R.K. (2016) The First Rule of Plant Transposable Element Silencing: Location, Location, Location. Plant Cell, 28, 304–313.

33. Araki, S., Le, N.T., Koizumi, K., Villar-Briones, A., Nonomura, K.-I., Endo, M., Inoue, H., Saze, H. and Komiya, R. (2020) miR2118-dependent U-rich phasiRNA production in rice anther wall development. Nat. Commun., 11, 3115.

34. Maguire, S., Lohman, G.J.S. and Guan, S. (2020) A low-bias and sensitive small RNA library preparation method using randomized splint ligation. Nucleic Acids Res., 10.1093/nar/gkaa480.

35. Lutzmayer, S., Enugutti, B. and Nodine, M.D. (2017) Novel small RNA spike-in oligonucleotides enable absolute normalization of small RNA-Seq data. Sci. Rep., 7, 5913.

